# Phosphoregulation of hierarchical protein condensate formation during clathrin-mediated endocytosis

**DOI:** 10.64898/2026.09.17.752504

**Authors:** Yidi Sun, Andrew Wong, Gean Hu, Albert Yeam, Katelyn Wong, Lucien Vu Dao, Aline Tschanz, David G. Drubin

## Abstract

The sequential assembly of over 50 proteins during clathrin-mediated endocytosis (CME) is well documented in yeast. However, the mechanisms governing this ordered recruitment remain poorly understood. Biomolecular condensation is proposed to orchestrate CME, but defining biophysical parameters *in vivo* remains challenging. Here, we developed a quantitative *in vivo* titration strategy to systematically determine CME protein assembly profiles. Beyond the previously reported Ede1 (yeast Eps15), seven additional proteins formed enlarged, concentration-dependent assemblies with distinct biophysical properties. Single-cell profiling revealed two biophysical classes among these eight proteins: homeostatic buffers with defined saturation boundaries and continuous assemblers that scale linearly. Acting as scaffolds and clients, these conserved proteins form an interconnected network of kinases, phosphatase recruiters, and substrates. Within this network, reversible phosphorylation modulates assembly stability to drive recruitment transitions. Collectively, this work establishes a two-stage model of protein condensation that drives CME progression from initiation to maturation and offers a general framework for how protein phosphorylation dynamically modulates biomolecular condensation *in vivo*.

**Short Summary:** Quantitative *in vivo* titration shows that eight yeast endocytic proteins form distinct, enlarged assemblies. Reversible phosphorylation dynamically tunes assembly thresholds, orchestrating a two-stage condensate reorganization from initiation to maturation during CME.

## Introduction

Clathrin-mediated endocytosis (CME) is a conserved eukaryotic process that dynamically regulates plasma membrane composition and selectively takes up molecules from the extracellular milieu (McMahon and Boucrot, 2011). In the budding yeast *Saccharomyces cerevisiae*, dozens of proteins assemble at endocytic sites in a predictable spatiotemporal order (Fig. 1 A) (Lu et al., 2016; Owen et al., 2025). Because these proteins are exceptionally well characterized in terms of dynamics, molecular function, and evolutionary conservation, budding yeast, with its simplicity and powerful genetics, provides an ideal system to uncover general CME principles across species (Johnson, 2024; Kaksonen and Roux, 2018). However, the physical mechanisms driving the self-organization of endocytic proteins into functional CME sites remain unclear.

**Figure 1.**
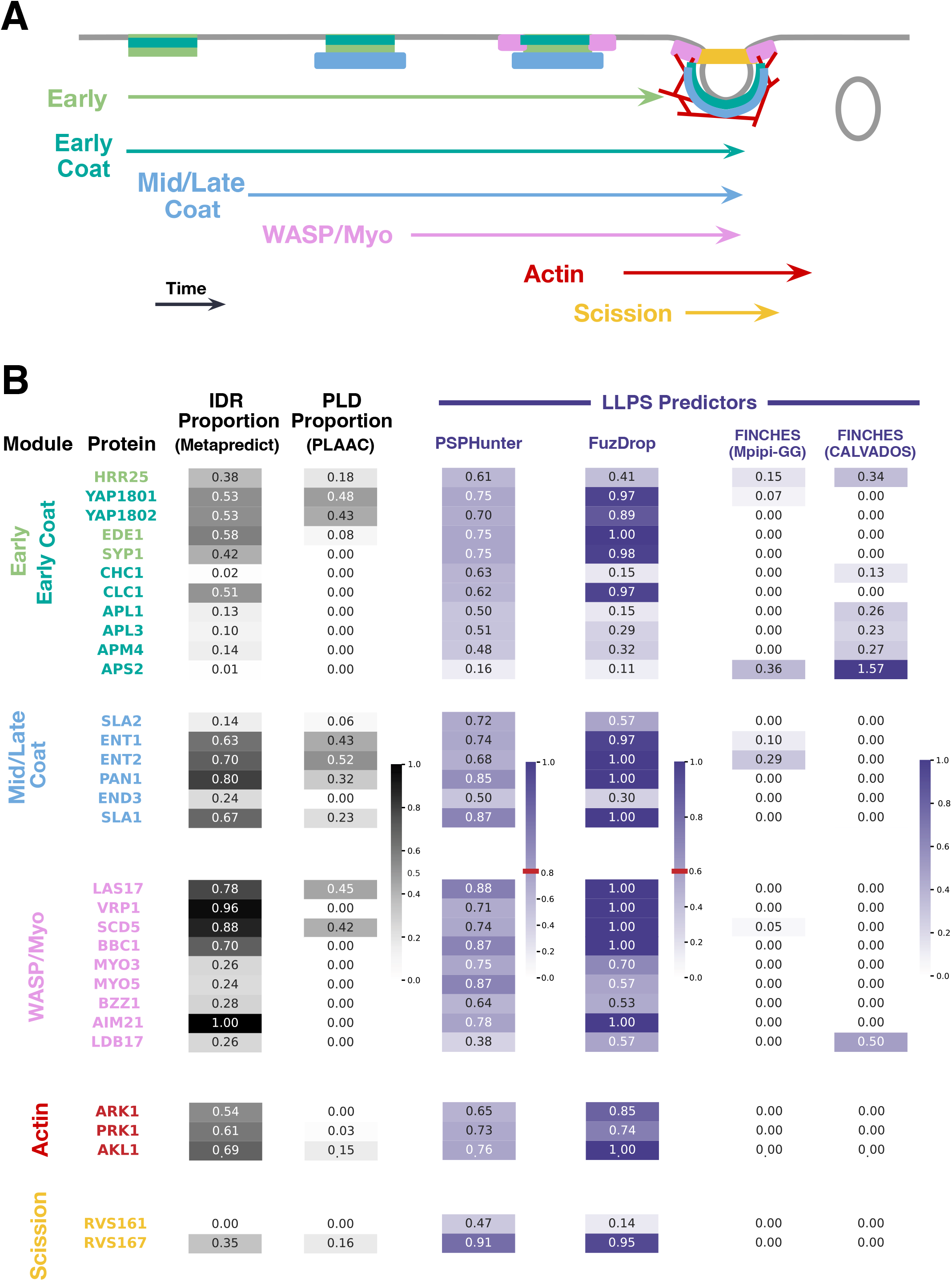
Spatiotemporal recruitment of yeast CME proteins and *in silico* CME protein condensation propensity predictions. **(A**) Schematic of the sequentially recruited yeast CME proteins and their organization into six modules. Site initiation begins with assembly of proteins in the early and early coat modules, followed by the mid/late coat module, then the WASp/myosin module to trigger local actin polymerization, and finally the scission module. Note that early modules, but not early coat modules, disassemble prior to robust actin assembly. **(B)** *In silico* analysis of phase separation propensity of the core yeast CME proteins, grouped by module. Heatmaps show predicted intrinsically disordered regions proportion (IDRs; Metapredict), prion-like domains proportion (PLDs; PLAAC), and liquid-liquid phase separation (LLPS) propensities using PSPHunter, FuzDrop, and FINCHES (employing the Mpipi-GG and the CALVADOS2 force field parameters, respectively). Red lines indicate classification thresholds. For the FINCHES prediction, the low-complexity region of the well-characterized LLPS-driving protein FUS was used as a positive control, with its propensity set as 1 (see Methods). Most actin-binding proteins (e.g., capping proteins Cap1/Cap2, the Arp2/3 complex) were excluded from this list.

Traditional mass-action models relying on stoichiometric protein-lipid and protein-protein interactions face biophysical constraints at the crowded CME site. First, low local concentrations of early initiation factors can delay site establishment, leaving strict 1:1 binding highly sensitive to cytoplasmic concentration fluctuations (Boeke et al., 2014; Klosin et al., 2020). Second, passive 3D cytoplasmic diffusion alone fails to rapidly concentrate endocytic machinery at the plasma membrane to meet demanding kinetic requirements (Day et al., 2021). Finally, overcoming high turgor pressure to drive membrane invagination in yeast requires that actin polymerization generate immense localized forces (Lacy et al., 2018; Lemiere and Chang, 2023; Nickaeen et al., 2019). Without a cohesive, cooperative network to distribute these loads, such forces risk rupturing individual molecular linkages or causing mechanical slippage across fluid lipid bilayers (Dmitrieff and Nedelec, 2015; Ren et al., 2023).

To overcome these thermodynamic and mechanical limits, biomolecular condensation offers an attractive organizing mechanism (Alberti et al., 2019; Banani et al., 2017; Choi et al., 2020; Deviri and Safran, 2021; Lyon et al., 2021; Mittag and Pappu, 2022). By forming mesoscale hubs, condensation could locally concentrate downstream adaptors to accelerate assembly kinetics. Indeed, mammalian initiators like Eps15 and Fcho1/2 co-assemble into liquid-like condensates acting as kinetic catalysts on membrane surfaces (Day et al., 2021), while yeast and plant homologs (Ede1 and AtEH proteins, respectively) similarly cluster early endocytic factors (Barkley et al., 2024; Kozak and Kaksonen, 2022). As the coat matures, this cohesive matrix is proposed to generate the interfacial tension required to bend the plasma membrane inward (Bergeron-Sandoval et al., 2021) and establish concentration thresholds that activate actin assembly (Sun et al., 2017). Although these models require further validation, such cross-kingdom examples implicate biomolecular condensation as a conserved principle of CME (Wilfling et al., 2023). Furthermore, because the CME network is transient, protein turnover must be dynamically regulated. While post-translational modifications (PTMs) such as phosphorylation are known to rewire binary coat protein interactions (Sekiya-Kawasaki et al., 2003; Zeng et al., 2007), whether and how phosphorylation modulates the material properties of CME condensates remains unexplored.

Elucidating the biophysical properties of protein assemblies *in vivo* is essential for understanding their functions in biomolecular condensation (Choi et al., 2020). In living cells, primary scaffolds act as homeostatic buffers, capping soluble cytosolic concentration (*C_L_*) at a defined buffering threshold (*C_onset_*) and routing excess mass into condensates (Alberti et al., 2019; Klosin et al., 2020). Conversely, weak assemblers and clients fail to buffer the cytosolic levels, causing *C_L_* to scale linearly with total protein concentration (*C_total_*). While classifying endocytic components into these regimes is critical to understand how CME assembly is driven, doing so remains challenging: native CME sites are sub-diffraction structures, whereas *in vitro* reconstitutions lack cellular crowding, network complexity, and proper PTM regulation (Banani et al., 2017; Riback et al., 2020). To address these challenges, we tested whether we could quantitatively analyze phase behavior *in vivo* by driving CME protein expression well beyond its cellular phase-separation threshold to generate optically resolvable assemblies.

Here, we first evaluated phase-separation propensities across the core yeast endocytic proteome using sequence-based algorithms, but mixed outputs did not narrow down specific candidates driving condensation. We then deployed a quantitative live-cell titration platform, leveraging progressive induction over time and natural single-cell expression variability, to systematically analyze phase behavior across 22 core CME proteins in yeast. Beyond Ede1, the early-arriving condensate-forming CME protein (Kozak and Kaksonen, 2022), our screen identified seven additional proteins recruited across distinct stages of CME that formed concentration-dependent enlarged assemblies *in vivo.* Single-cell concentration profiling (*C_L_* vs. *C_total_*) partitioned these eight assembly-forming proteins into two discrete biophysical classes: robust homeostatic buffers with operational onset thresholds (*C_onset_*), and continuous, low-affinity homotypic assemblers. Intriguingly, these eight components form an interconnected kinase-phosphatase circuit. Reversible phosphorylation actively modulates *C_onset_*, driving a coordinated two-stage condensation transition from CME site initiation to maturation. Given the evolutionary conservation of the CME machinery, this quantitative *in vivo* framework establishes broadly applicable principles for how protein phosphorylation dynamically tunes biomolecular phase boundaries to drive CME site assembly and turnover.

## Results

### *In silico* profiling of endocytic protein condensation propensity

To predict which CME proteins across all six functional CME modules are most likely to drive condensate formation, we screened core endocytic proteins using computational predictors of phase separation propensity and related sequence features (Fig. 1 B) (See Methods).

Intrinsically disordered regions (IDRs) can drive biomolecular condensation when they feature specific sequence grammars, such as prion-like domains (PLDs) (Mittag and Pappu, 2022). While most CME proteins analyzed contain high predicted IDR content (metapredict; Emenecker et al., 2021), only a subset harbors PLDs (PLAAC; Lancaster et al., 2014) (Fig. 1 B). Several of these PLD-containing CME proteins, such as Yap1801/2 (yeast CALM/AP180), Ent1/2 (yeast Epsin), and Sla1 (yeast CIN85), have been proposed to form endocytic condensates (Bergeron-Sandoval et al., 2021).

Using machine-learning-based predictors, we evaluated LLPS propensity (FuzDrop; Hardenberg et al., 2020; PSPHunter; Sun et al., 2024a). High LLPS propensity was predicted for most CME proteins (Figure 1 B). However, these data-driven models may be biased as their training sets include several CME factors (see Methods). In contrast, FINCHES, a physics-driven model of IDR interactions (Ginell et al., 2025), predicted few proteins with high LLPS propensity. These ambiguous *in silico* predictions render a systematic live-cell experimental strategy essential to understand phase separation in CME.

### An *in vivo* platform to quantitatively evaluate condensate-forming candidates

Because biomolecular condensation inherently depends on surpassing a critical saturation threshold *C_sat_*, evaluating phase behavior *in vivo* requires titrating protein expression across a broad concentration range. To achieve this, we established a quantitative live-cell titration platform (Fig. 2). We varied GFP-tagged CME protein levels by replacing the native promoters of their genes with a *β*-estradiol–inducible *GAL1* promoter driven by Gal4-ER-VP16 (GEV) (Fig. 2 A) (McIsaac et al., 2011). This strategy enables rapid expression induction without switching yeast media from glucose to galactose, avoiding confounding metabolic alterations. Control experiments confirmed that *β*-estradiol exposure alone did not alter wild-type CME dynamics (Fig. S1 A). By progressively increasing expression beyond endogenous levels, this platform enabled us to screen for condensate-forming candidates, which appeared as enlarged puncta distinct from native CME sites in size and brightness. Moreover, low basal expression in the uninduced state maintained sub-physiological levels for essential proteins sufficient to support cell viability. Induction levels, dynamic range, and full-length fusion protein integrity were confirmed by quantitative Western blot analysis.

**Figure 2.**
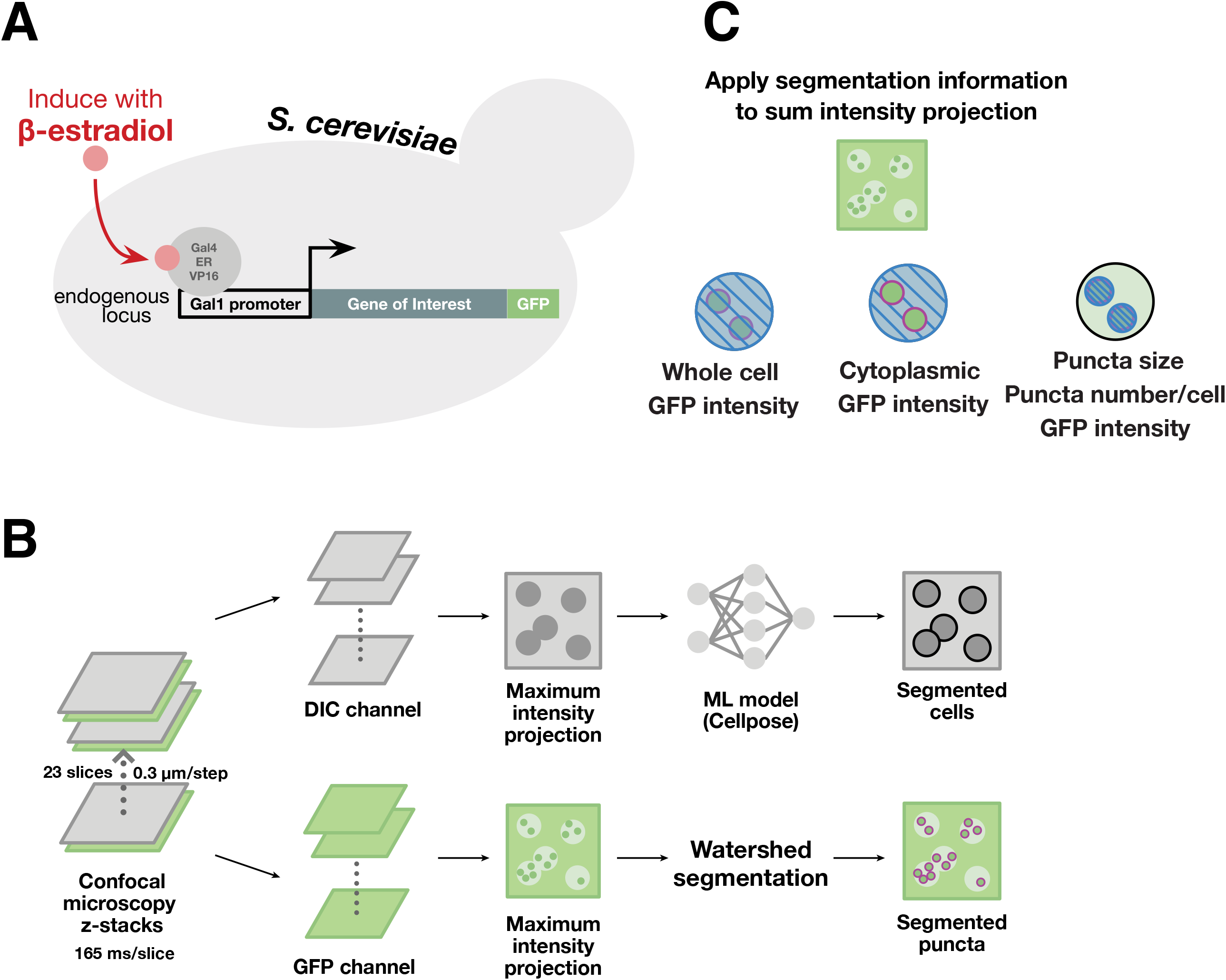
Inducible gene overexpression and automated quantitative image analysis of enlarged concentration-dependent assemblies. **(A)** Schematic of the *β*-estradiol–inducible system used to tune overexpression of GFP-tagged proteins expressed from endogenous loci, with expression levels assessed by Western blot. **(B)** Automated processing pipeline for 3D confocal z-stacks. Cell boundaries are segmented from DIC maximum intensity projections (MIPs) using Cellpose, while puncta are segmented from GFP-channel MIPs using watershed algorithms. **(C)** Extracted single-cell and puncta quantification metrics from GFP sum intensity projections (SIPs). See Methods for details.

To rigorously quantify candidate condensates identified during visual screening, we acquired z-stack confocal data, collecting 23 slices at a 0.3 *μ*m step size and at an acquisition rate of 165 ms per slice to measure total intracellular GFP signals (Fig. 2 B). Cell segmentation was performed using Cellpose (Stringer et al., 2021), and a custom pipeline isolated data for individual cells and their intracellular puncta (Fig. 2 B). From each segmented cell, the algorithm extracted the mean GFP intensity (*I_total_)* from the sum intensity projections (Fig. 2 C). The custom code then masked identified puncta to exclude their signals and measure the median intensity of the dilute cytoplasm (*I_L_*) for each cell (Fig. 2 C) (see Methods). Puncta size, frequency, and mean assembly intensity (*I_D_*, representing the puncta intensity) were also quantified (Fig. 2 C).

To distinguish the overexpression-induced foci from faint, diffraction-limited native CME sites without presupposing phase separation, we refer to these foci observationally as enlarged puncta and molecularly as assemblies. Combining tunable gene expression with automated image analysis provides a quantitative platform to evaluate concentration-dependent assembly formation.

### Establishing and benchmarking an *in vivo* quantitative platform via overexpression-induced Ede1 enlarged puncta formation

To validate our approach, we analyzed Ede1, an early-arriving endocytic protein known to form enlarged puncta upon overexpression (Kozak and Kaksonen, 2022; Wilfling et al., 2020). Upon *β* -estradiol induction of a *GAL1-EDE1-GFP* strain, we tracked Ede1-GFP expression kinetics and intracellular behavior over a 180-min time-course (Fig. 3). Endogenous Ede1-GFP expression driven by native *EDE1* promoter served as the baseline control.

**Figure 3.**
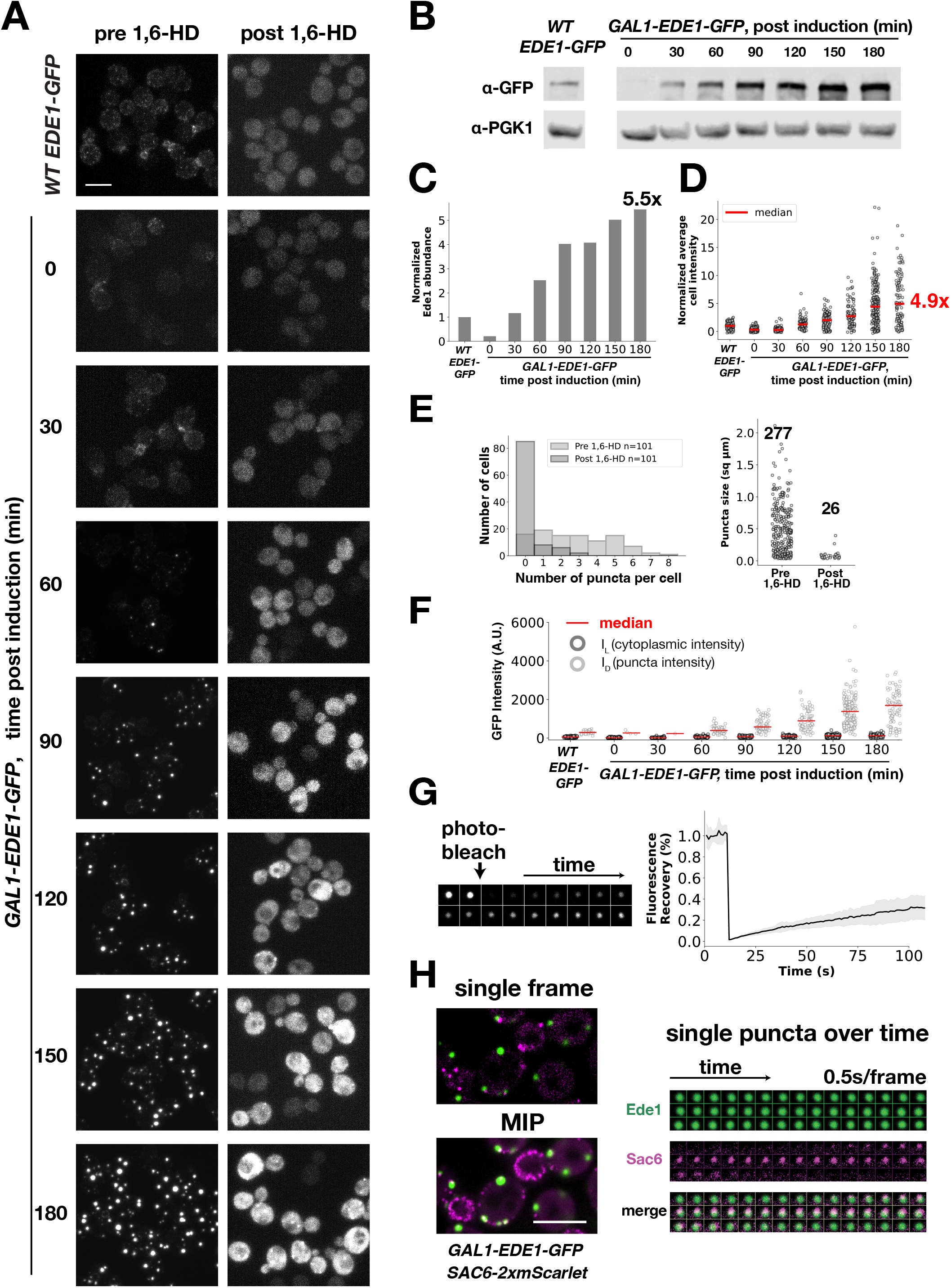
Benchmarking the dose-response platform using *Ede1*. **(A)** MIPs of z-stack live cell imaging before (left) and after (right) 5 min treatment with 10% 1,6-hexanediol (1,6-HD). Top row: wild-type (*WT*) cells expressing Ede1-GFP from its endogenous promoter. Lower rows: *GAL1-EDE1-GFP* cells induced with 50 nM *β* -estradiol across the indicated time points. **(B, C)** Western blot **(B)** and Pgk1-normalized quantification **(C)** of Ede1-GFP abundance relative to wild-type across the induction time course. **(D)** Total cellular fluorescence intensity of Ede1-GFP relative to wild-type. Each circle represents an individual cell; red bars indicate medians. **(E)** Puncta frequency per cell (left) and puncta size (right) pre-and post-1,6-HD treatment (*n* = 101 cells per group). **(F)** Dilute cytoplasmic (*I_L_*) and dense puncta (*I_D_*) fluorescence intensities across the induction. **(G)** Representative FRAP montage of a single punctum (left) and mean fluorescence recovery curve ± S.D. (right). **(H)** Two-color live-cell imaging of Ede1-GFP and Sac6-2xmScarlet. Left to right: single mid-focal plane frame, MIP from a 2 min movie, and time-lapse montage of a single punctum. The MIP represents all the native CME events that occurred alongside concentration-dependent Ede1-GFP puncta assembled over the course of the movie. Scale bar, 5 *μ*m

Under uninduced conditions (*t* = 0), Ede1-GFP forms faint cortical patches corresponding to CME sites (Fig. 3 A). Within 60 min post-induction, bright, enlarged puncta emerged at the plasma membrane and in the cytoplasm, progressively growing in size and intensity (Fig. 3 A). Quantitative Western blot analysis confirmed expression of full-length Ede1-GFP, with total protein levels reaching 5.5-fold over WT at 180 min (Fig. 3, B and C). Single-cell imaging mirrored this increase, showing a steady rise in total fluorescence intensity (*I_total_*) to a median of 4.9-fold over baseline (Fig. 3 D), with individual cells reaching up to 22.2-fold (Table S1).

We then evaluated the biophysical properties of these induced Ede1 assemblies. First, we used a 1,6-hexanediol (1,6-HD) assay to probe assembly stability and sensitivity to hydrophobic disruption (Alberti et al., 2025; Duster et al., 2021). A brief 5-min 1,6-HD treatment completely dissolved puncta induced for less than 90 min, while puncta induced for 180 min were reduced in frequency by 90.6% with sharply decreased average size (Fig. 3, A and E). Second, we quantified Ede1-GFP partitioning between the dilute cytoplasm (*I_L_*) and dense assemblies (*I_D_*) across the induction time course. As total intensity (*I_total_*) rose (Fig. 3 D), *I_L_*remained low and steady, whereas *I_D_*increased substantially (Fig. 3 F). This asymmetric partitioning indicates that the cytosolic phase concentration is buffered, with excess protein routed into dense assemblies, a hallmark of concentration-dependent cellular protein condensation (Klosin et al., 2020; Riback et al., 2020). Fluorescence recovery after photobleaching (FRAP) (Phair et al., 2004) confirmed dynamic molecular exchange between Ede1 assemblies and the cytoplasm (Fig. 3 G). Fitting a single-exponential model to recovery curves (n = 9) yielded a half-time (*t*_1/2_) of 74 s and a mobile fraction (*M_f_*) of 55%.

Fast two-color live-cell imaging (0.5 s/frame) at a single equatorial focal plane was performed by image scanning microscopy (AXR with Nikon NSPARC). The enlarged Ede1-GFP puncta were primarily cortical and stationary, demonstrated by near-perfect overlap between individual frames and the maximum intensity projection (MIP) across a 2 min movie (Fig. 3 H). At these stationary Ede1 assemblies, the actin-bundling protein Sac6 (yeast fimbrin), dynamically appeared and disappeared (Fig. 3 H), confirming the functional engagement with CME actin assembly machinery. Endogenous CME persisted alongside these induced assemblies, as evidenced by polarized cortical Sac6 foci in the MIPs (Fig. 3 H).

Together, these benchmark results with Ede1 validated that our dose-response platform effectively induces assembly formation. Integrating 1,6-HD sensitivity, cytosolic buffering and FRAP analysis, and functional actin assembly assays establishes a comprehensive pipeline to evaluate the biophysical properties of candidate condensate-forming CME proteins.

### A systematic screen of twenty-one core CME proteins identifies seven additional proteins undergoing concentration-dependent assembly upon overexpression

To systematically survey the propensity of endocytic proteins to undergo concentration-dependent assembly *in vivo*, we constructed an inducible-expression library for 21 core CME proteins (Figs. 1 B and 2 A; and Table S1). We excluded multisubunit complexes requiring tightly coordinated stoichiometric expression, such as clathrin and the AP-2 adaptor complex, from the screen.

Following a 180 min *β*-estradiol induction, confocal live-cell imaging revealed that 14 of the 21 candidates failed to form enlarged, visually apparent puncta (Fig. S2). These non-hits included major adaptors (such as Yap1801/2 and Ent1/2) and key actin regulators (such as the yeast WASp, Las17, and type I myosin, Myo5). Protein expression level increases were verified by quantitative Western blotting and single-cell GFP intensity tracking (Fig. S2). Quantifying maximum single-cell fold changes relative to WT baselines confirmed high induction levels, ranging from 12.3-fold for Rvs167 (yeast amphiphysin) to 108-fold for Ldb17 (Table S1). Despite highly elevated cellular concentrations, these proteins retained relatively normal cortical patch localization.

Importantly, seven proteins from this screen, Pan1, Scd5, Sla2, Hrr25, Akl1, Sla1, and Bbc1, formed enlarged puncta upon induction (Figs. 4 and 5; Fig. S1 B; Table S1). Notably, Pan1 (yeast intersectin ITSN) and Scd5 (PP1 phosphatase-targeting subunit) are essential for vegetative growth. Pan1, a key scaffold of the CME protein network (Bradford et al., 2015; Miliaras et al., 2004; Sun et al., 2015), is a substrate for the phosphatase recruited by Scd5 (Zeng et al., 2007). Sla2 (yeast Hip1R and Hip1) links the endocytic coat to the polymerizing actin network, transmitting mechanical force during membrane invagination (Kaksonen et al., 2003; Skruzny et al., 2012). Two protein kinases were also identified: Hrr25 (yeast casein kinase 1), which phosphorylates Ede1 to promote site initiation (Peng et al., 2015), and Akl1 (Ark1/Prk1 family, yeast AAK1/GAK), which phosphorylates coat components to promote CME site disassembly (Bourgoint et al., 2018; Huang et al., 2003; Roelants et al., 2017; Sekiya-Kawasaki et al., 2003). Finally, Sla1 (yeast CIN85) and Bbc1 act as critical regulators of Las17-mediated actin nucleation through the SH3 (Src Homology 3)-PRD (Proline-Rich Domain) network (Feliciano and Di Pietro, 2012; Rodal et al., 2003). Direct visual comparison of these new candidates alongside Ede1 revealed distinct puncta morphologies, frequencies, and cytosolic fluorescence intensities (Fig. S1 B). Thus, by capturing key components across the endocytic pathway, our screen suggests that biomolecular condensation may function beyond CME initiation. Next, we applied the Ede1-benchmarked pipeline (Fig. 3) to systematically evaluate these candidate assemblies.

**Figure 4.**
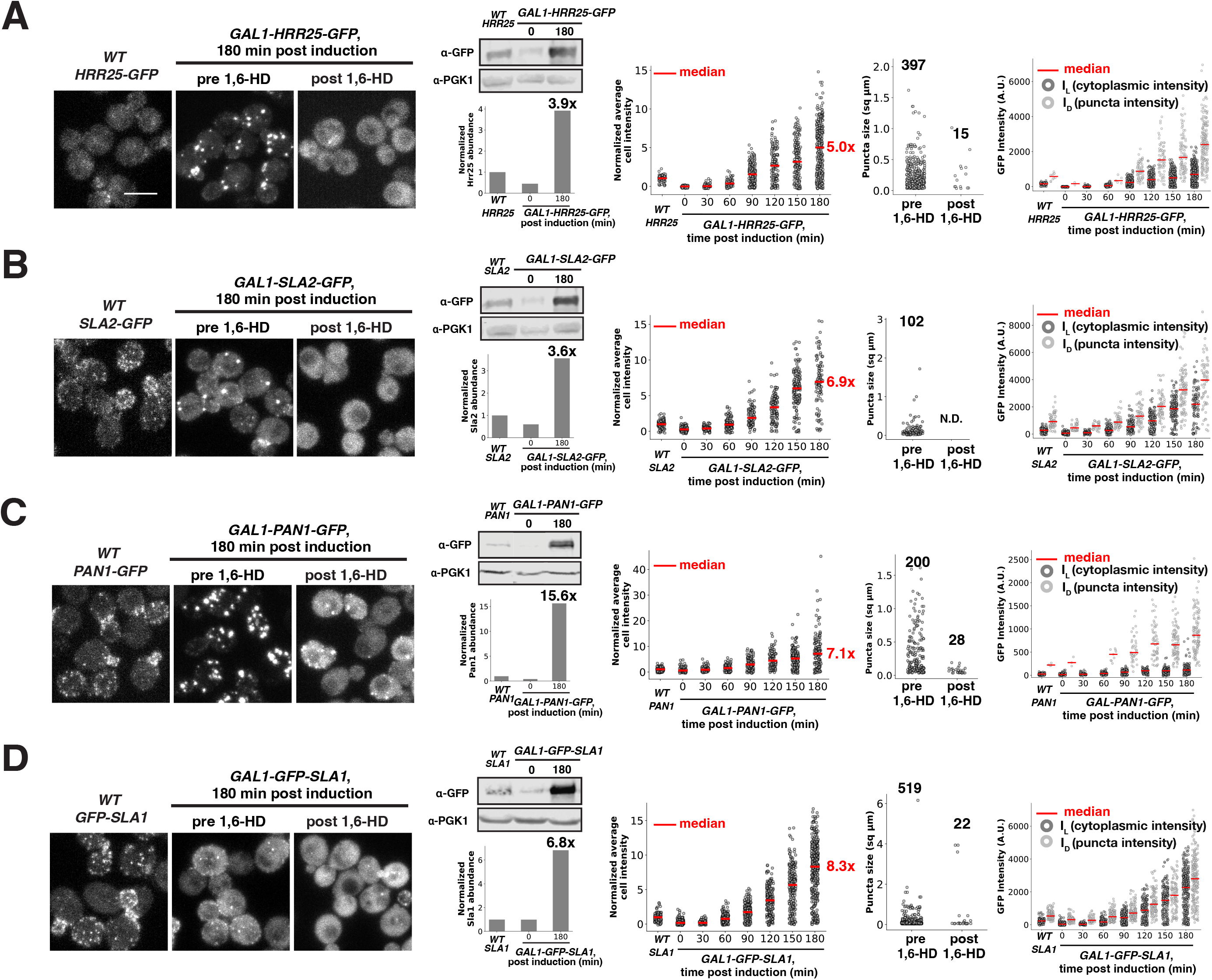
1,6-HD sensitivity assays and intracellular partitioning of enlarged concentration-dependent assemblies of early- and middle-arriving CME proteins (A–D) Analysis of overexpressed GFP-tagged CME proteins: **(A)** Hrr25, **(B)** Sla2, **(C)** Pan1, and **(D)** Sla1. For each protein, panels from left to right: MIPs from z-stack live-cell imaging of wild-type cells endogenously expressing indicated GFP tagged protein, and cells expressing the *GAL1*-driven GFP-tagged protein following 180 min *β* -estradiol induction before and after 5 min treatment with 10% 1,6-HD; Western blot and Pgk1-normalized quantification of protein abundance at 0 and 180 min post-induction relative to wild-type; total cellular fluorescence intensity relative to wild-type across the induction time course; puncta size (*μ*m^2^) and number before and after 1,6-HD treatment (values above scatter plots indicate the number of puncta detected; N.D., not detected); and *I_L_* and *I_D_* across the induction time course. Scale bar, 5 *μ*m

**Figure 5.**
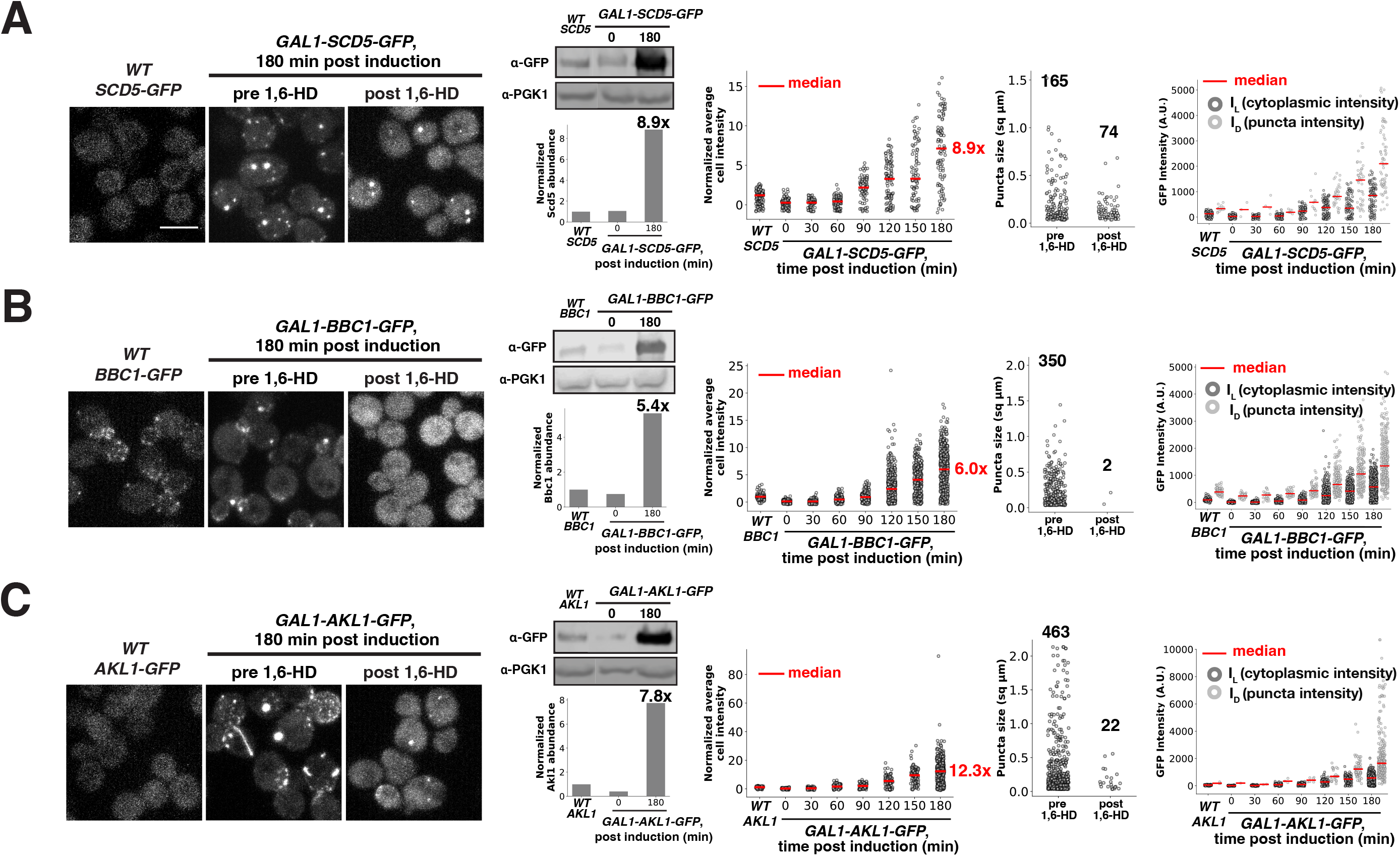
1,6-HD sensitivity assays and intracellular partitioning of enlarged concentration-dependent assemblies of three late-arriving CME proteins (A–C) Analysis of overexpressed GFP-tagged CME proteins: **(A)** Scd5, **(B)** Bbc1, and **(C)** Akl1. For each protein, panels from left to right: MIPs from z-stack live cell imaging of wild-type cells endogenously expressing indicated GFP tagged protein, and cells expressing the *GAL1*-driven GFP-tagged protein following 180 min *β*-estradiol induction before and after 5 min treatment with 10% 1,6-HD; Western blot and Pgk1-normalized quantification of protein abundance at 0 and 180 min post-induction relative to wild-type; total cellular fluorescence intensity relative to wild-type across the induction time course; puncta size (*μ*m^2^) and number before and after 1,6-HD treatment (values above scatter plots indicate the number of puncta detected); and *I_L_* and *I_D_*across the induction time course. Scale bar, 5 *μ*m

### 1,6-hexanediol treatment reveals distinct physical properties across candidate assemblies

Quantitative Western blotting and single-cell fluorescence quantification confirmed robust, full-length expression of each candidate protein relative to WT after 180 min induction (Figs. 4 and 5). Although both methods showed consistent trends, their absolute quantitative values did not always directly align. This divergence most likely reflects cell-to-cell heterogeneity in induction, as bulk Western blotting yields population-wide averages, whereas single-cell fluorescence quantification reports individual cell medians. Pan1-GFP exemplified this distinction: the population-wide Western blot showed a 15.6-fold increase, whereas its median single-cell intensity increased by 7.1-fold (Fig. 4 C). Single-cell tracking captured maximum Pan1 expression levels of up to 45.1-fold within individual cells (Table S1), highlighting a right-skewed population where high-expressing outliers disproportionately shift the pooled biochemical average.

As with Ede1, we used 1,6-HD treatment to evaluate the relative contribution of weak hydrophobic interactions across these candidate assemblies (Figs. 4 and 5). Hrr25, Sla2, Sla1, and Bbc1 exhibited pronounced sensitivity to 1,6-HD, with puncta frequency decreasing by 95%-100% and remaining assemblies shrinking noticeably (Figs. 4 A, B, and D; Fig. 5 B; and Table S1). Uniquely, overexpressed Akl1 generated two morphologically distinct populations of assemblies: cortical enlarged puncta and elongated, cable-like structures (Fig. 5 C). 1,6-HD treatment dissolved most Akl1 assemblies, reducing the overall frequency by 95.3%. In contrast, Pan1 assemblies were moderately resistant (86% reduction; Fig. 4 B), whereas Scd5 assemblies proved the most resistant, showing only a 55% drop in frequency with largely unaffected size distributions (Fig. 5 A). This differential susceptibility suggests that while weak hydrophobic forces drive assembly across all candidates, Pan1 and Scd5 form more resistant structures, potentially stabilized by higher network valency, electrostatic interactions, or distinct material states.

### Partitioning of candidate proteins between the cytoplasm and dense assemblies reveals a spectrum of buffering behaviors

To determine whether the seven new candidates exhibit cytosolic buffering, we monitored protein partitioning between the dilute cytoplasm (*I_L_*) and dense assemblies (*I_D_*) over a 180 min induction time course (Figs. 4 and 5). Individual tracking profiles revealed distinct partitioning trends between *I_L_* and *I_D_*across varying total cellular expression levels. For a subset of candidates, including Hrr25, Sla2, Sla1, and Bbc1, the intensities of both *I_L_* and *I_D_* scaled proportionally over time, showing no apparent plateau in the cytoplasmic signal (Figs. 4, A, B, and D; 5 B). Thus, as total protein abundance increases, these proteins continue to partition into cytosolic and dense fractions without cytoplasmic buffering. In contrast, Pan1 and Akl1 showed distinct biphasic behavior: *I_L_* initially rose (0-120 min) before flattening as *I_D_* continued to accumulate, reflecting cytosolic buffering at higher expression levels (Figs. 4 C and 5 C). Finally, Scd5 exhibited a unique biphasic profile, where *I_L_* displayed transient sub-linear buffering at intermediate induction times (90-150 min) before rising concurrently with *I_D_* at 180 min, while maintaining an overall attenuated accumulation relative to total protein levels (Fig. 5 A). Together, these results uncover distinct regimes of concentration buffering among the seven CME candidates.

### Candidate assemblies differentially recruit actin assembly machinery and exhibit diverse molecular dynamics

Next, to evaluate protein dynamics in the dense assemblies and their ability to recruit downstream actin assembly machinery, we performed two-color live-cell equatorial imaging with the actin reporter Sac6 after 180 min of induction (Figs. 6 and 7). Hrr25, Sla2, and Bbc1 form stationary enlarged assemblies at the cell cortex upon induction, as evidenced by near-perfect overlap between individual frames and their MIP images from 2-min movies (Fig. 6, A-C). Sac6 transiently assembled and disassembled at these structures, suggesting they retain the ability to recruit downstream actin assembly machinery. In contrast, Sac6 was undetectable at the spherical or cable-like stationary Akl1 assemblies (Fig. 6 D). Cortical Sac6 foci not associated with the enlarged assemblies represent native CME sites. While these native sites remained intact upon induction of Hrr25, Sla2, and Bbc1 overexpression, MIP images showed that cortical Sac6 foci were severely diminished in Akl1-overexpressing cells (Fig. 6). These results suggest that excessive Akl1 disrupts actin assembly even at native CME sites.

**Figure 6.**
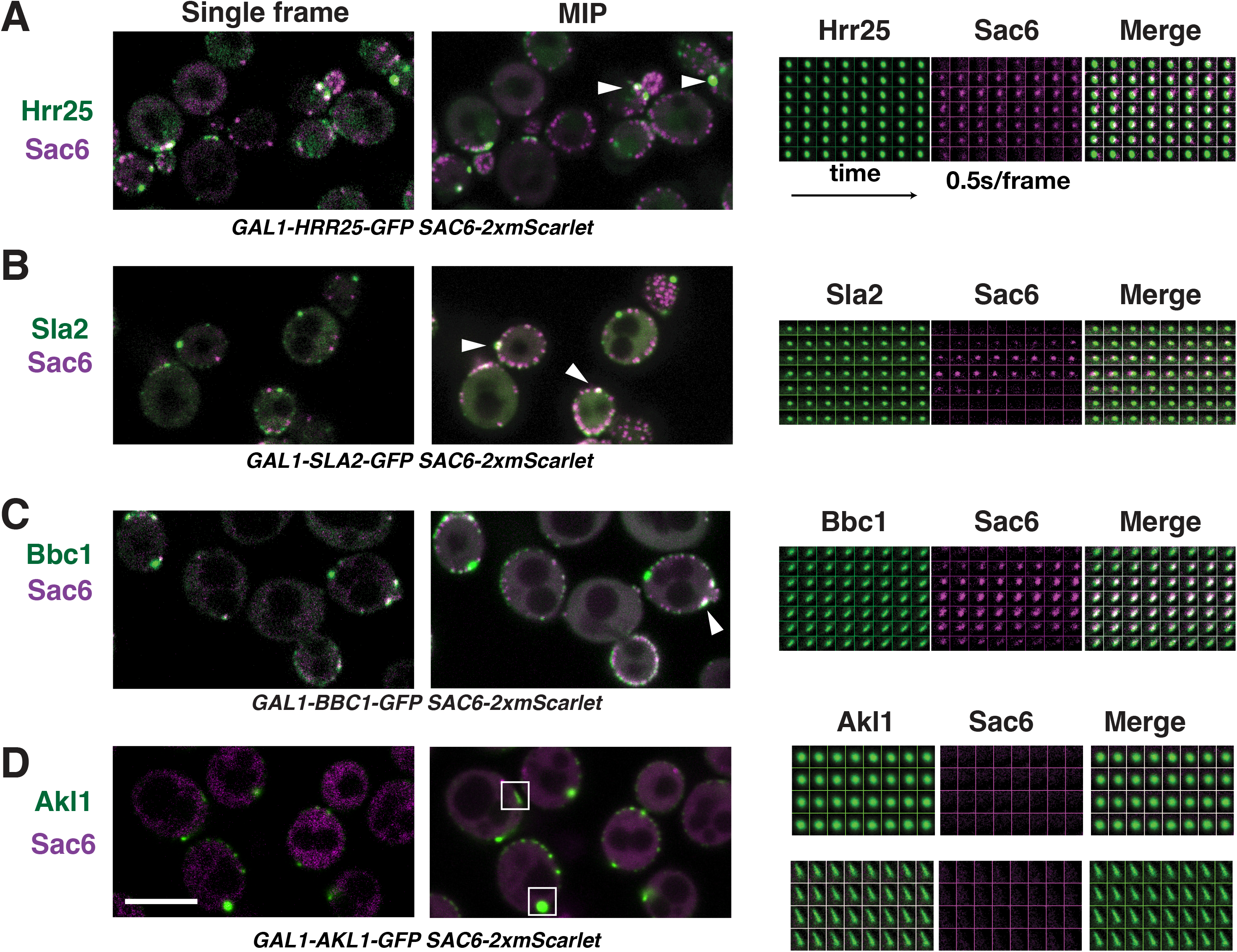
Actin assembly at stationary enlarged concentration-dependent assemblies of CME proteins (A–D) Two-color live-cell imaging of GFP-tagged proteins (**A**, Hrr25; **B**, Sla2; **C**, Bbc1; **D**, Akl1) and Sac6-2xmScarlet following 180 min of *β*-estradiol induction. For each protein, panels from left to right show: a single mid-focal plane frame, a MIP of the 2 min movie, and a time-lapse montage of a single punctum (0.5 s/frame). Panel D includes two representative montages. Arrows mark the representative stationary puncta colocalized with Sac6. Boxes indicate Akl1 puncta and cable-like structures lacking Sac6. Scale bar, 5 *μ*m

**Figure 7.**
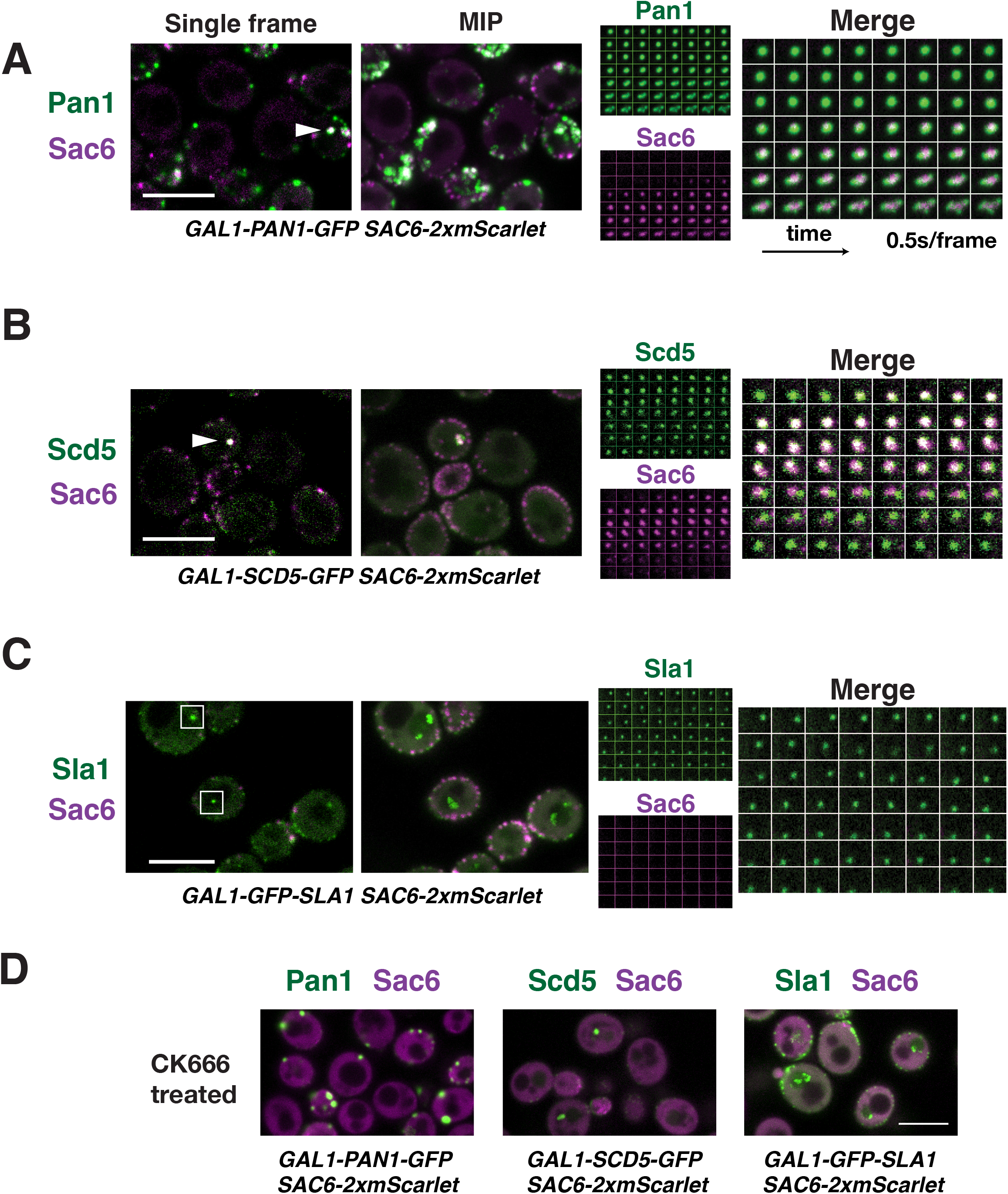
Actin assembly at motile enlarged concentration-dependent assemblies of CME proteins (A–C) Two-color live-cell imaging of GFP-tagged proteins (**A**, Pan1; **B**, Scd5; **C**, Sla1) and Sac6-2xmScarlet following 180 min of *β*-estradiol induction. For each protein, panels from left to right show: a single mid-focal plane frame, a MIP of the 2 min movie, and a time-lapse montage of a single punctum (0.5 s/frame). Arrows mark the representative stationary puncta colocalized with Sac6. Boxes indicate Sla1 puncta lacking Sac6. **(D)** Two-color live-cell imaging of GFP-tagged proteins (left, Pan1; middle, Scd5; right, Sla1) and Sac6-2xmScarlet following 180 min of *β*-estradiol induction and subsequent treatment with 250µM CK666 for 5-10 min. Note the diffuse cytoplasmic localization of Sac6, indicating successful Arp2/3 inhibition and consequent actin disassembly. Scale bar, 5 *μ*m

In contrast, Pan1 formed highly dynamic assemblies at the cell cortex and in the cytoplasm upon overexpression (Fig. 7 A). The MIP image from a 2-min movie appeared smeared compared with a single time frame, reflecting the dynamic nature of Pan1 assemblies. These Pan1 assemblies initially appeared spherical, reminiscent of phase-separated droplets, and subsequently recruited Sac6. It appeared that the resulting actin assembly forces pushed and deformed the Pan1 assemblies, splitting them into several smaller assemblies as Sac6 disassembled (Fig. 7 A). To test whether local Arp2/3-mediated actin polymerization drove this assembly remodeling, cells were treated with the Arp2/3 inhibitor CK-666 (Hetrick et al., 2013). This treatment completely blocked actin assembly and Pan1 assembly motility and deformation, maintaining Pan1 assemblies as intact spherical droplets (Fig. 7 D). Overexpressed Scd5 formed fewer and smaller cytoplasmic assemblies that nonetheless exhibited actin assembly and Arp2/3-dependent motility like with Pan1 (Figs. 7 A, B, and D). In contrast, Sac6 was undetectable at small cytoplasmic Sla1 assemblies despite their motility (Fig. 7 C), and Sla1 motility was insensitive to CK-666 treatment (Fig. 7 D). Thus, Sla1 assemblies lack the functional interactions required to trigger CME-associated actin assembly.

FRAP analysis was performed on the candidate assemblies to generate recovery curves evaluating exchange dynamics with the cytoplasmic pool, which were fitted to a single-exponential model (Fig. 8, A-F, and Table S1). Hrr25 assemblies exhibited moderate exchange kinetics (*t*_1/2_ = 23.4 s, *M_f_* = 39%) (Fig. 8 A), whereas Sla2 and Bbc1 displayed rapid exchange kinetics (*t*_1/2_ = 6.6 s and 7.9 s; *M_f_* = 35% and 55%, respectively) (Fig. 8, B and C). Because Pan1 and Scd5 assemblies are motile within cells, we immobilized them with CK-666 before photobleaching. Pan1 assemblies displayed relatively slow exchange kinetics ( *t*_1/2_ = 193.4 s , *M_f_* = 41%), whereas Scd5 exhibited relatively rapid recovery (*t*_1/2_ = 5.3 s, *M_f_* = 54%) (Fig. 8, D and E). Conversely, we could not perform FRAP analysis on Sla1 assemblies because of their persistent, CK-666-insensitive motility (Fig. 7, C and D). For Akl1, FRAP distinguished the biophysical properties of its two co-existing structures: spherical assemblies dynamically exchanged Akl1 with the cytoplasm ( *t*_1/2_ = 26 s , *M_f_* = 49%), whereas cable-like structures exhibited negligible recovery (Fig. 8 F). Collectively, these distinct recovery profiles show that molecular exchange kinetics between dense assemblies and the cytoplasm vary widely across these candidate CME proteins.

**Figure 8.**
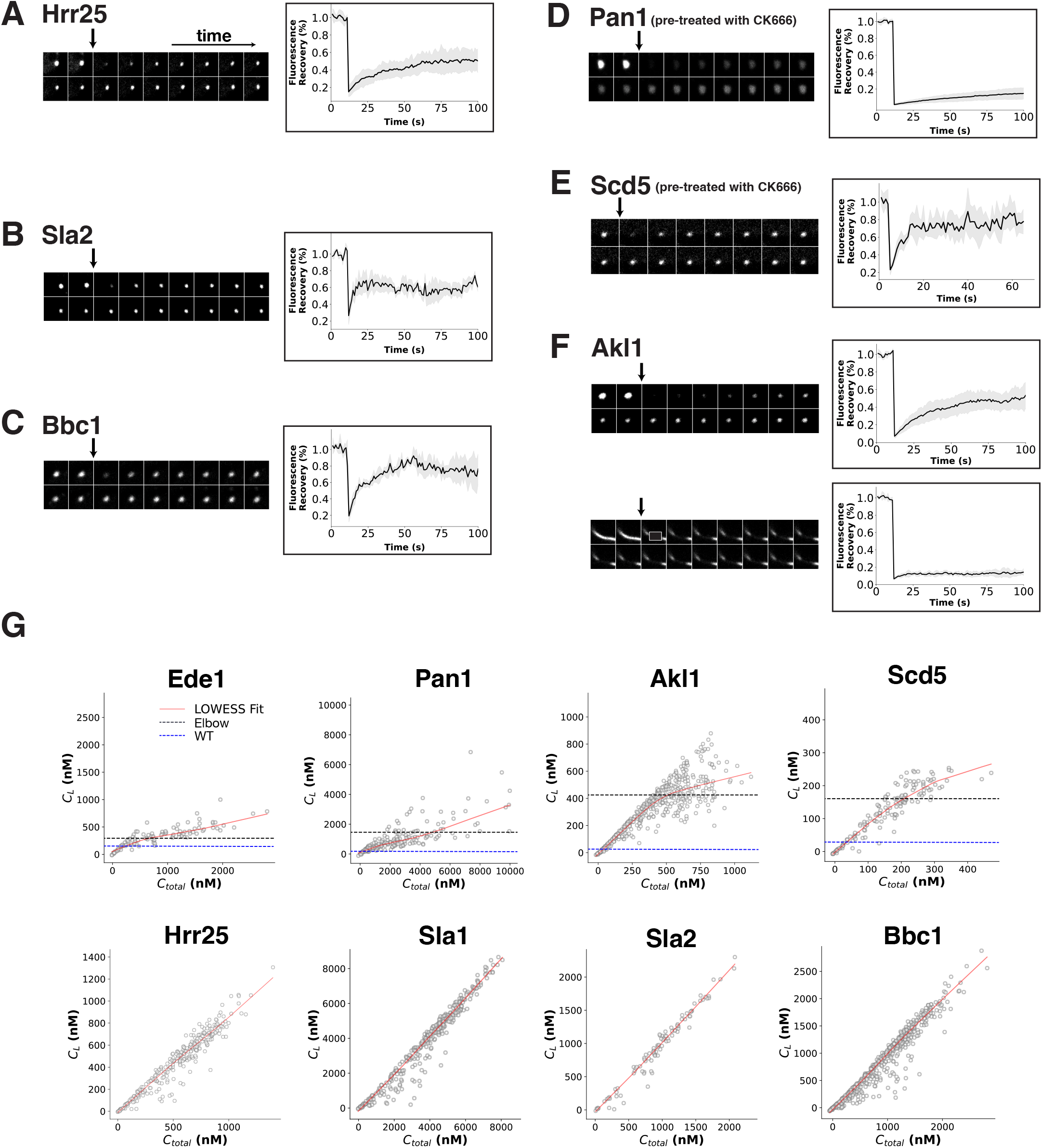
Dynamic exchange of enlarged concentration-dependent assemblies and single-cell dose-response profiling of CME proteins (A–F) Representative time-lapse montages (left) and mean fluorescence recovery after photobleaching (FRAP) curves (±S.D., right) for the indicated GFP-tagged proteins following 180 min of *β*-estradiol induction. Black arrows mark the photobleaching event. Table S1 summarizes the recovery half-time (*t*_1/2_), mobile fraction (*M_f_*), and sample size (*n*). Note that in the Pan1 **(D)** and Scd5 **(E)** FRAP studies, cells were treated with CK666 prior to photobleaching; and panel **(F)** shows representative FRAP montages and recovery curves for spherical (top) and cable-like (bottom) Akl1 structures. **(G)** Phase-separation partitioning diagrams showing dilute cytoplasmic concentration (*C_L_*) as a function of total cellular concentration (*C_total_*) for indicated proteins at 180 min induction time point. Each circle represents an individual cell. Red lines represent LOWESS fits, dark gray dashed lines indicate the estimated *C_onset_* operationally calculated as the elbow point along the LOWESS fit, and blue dashed lines indicate baseline wild-type (WT) cytoplasmic concentrations.

### Single-cell dose-response profiling classifies candidate assemblies into distinct partitioning regimes

Although tracking fluorescence intensities (*I_L_* vs *I_D_*) over time provided kinetic insights into assembly growth (Figs. 4 and 5), quantifying concentration buffering necessitated analyzing the pseudo-phase behavior of Ede1 and the seven new candidates *in vivo*. Generating *in vivo* pseudo-phase diagrams (*C_L_* vs *C_total_*) requires measuring fluorescence intensities across a wide expression range and converting them into absolute molar concentrations (Klosin et al., 2020; Riback et al., 2020). By leveraging natural cell-to-cell expression variability at 180 min and calibrating against protein abundances in wild-type cells (Boeke et al., 2014) (see Methods), we treated each cell as an individual titration point. For motile assemblies (Pan1 and Scd5), we used CK-666 to arrest motility prior to imaging. Ultimately, our results suggest a clear split among the eight proteins, distinguishing buffered components with distinct operational buffering onsets ( *C_onset_*) from unbuffered components (Fig. 8 G).

The first group, comprising Ede1, Pan1, Akl1, and Scd5, exhibited clear concentration-buffering behavior (Fig. 8 G). For these proteins, a smoothed trendline fitted to the single-cell data (LOWESS fit, see Methods) revealed that cytoplasmic concentration ( *C_L_)* initially scales proportionally with total cellular protein (*C_total_*). At higher expression levels, this trendline bends downward and levels off, defining an operational buffering onset (*C_onset_*) at the elbow point where soluble protein accumulation flattens into a sub-linear regime. This partial slope attenuation reflects dynamic cellular buffering in a non-equilibrium *in vivo* environment, where active cellular processes prevent the strict zero-slope plateau typical of simple *in vitro* equilibrium systems (Klosin et al., 2020; McSwiggen et al., 2019; Riback et al., 2020). Comparing these empirical thresholds (*C_onset_*) to wild-type levels (Boeke et al., 2014) revealed distinct buffering ranges. Ede1 began buffering at ∼284 nM (2-fold over its 138 nM wild-type baseline), whereas Pan1 reached a much higher threshold of ∼1357 nM (10-fold over its 140 nM baseline). Scd5 and Akl1 displayed thresholds of ∼161 nM and ∼412 nM, respectively (∼5-fold and 12-fold over their wild-type levels of 31 nM and 34 nM). Thus, all four proteins limit excess cytoplasmic buildup through concentration-dependent buffering.

In contrast, the second group, comprising Hrr25, Sla1, Sla2, and Bbc1, showed no evidence of a buffering threshold or slope attenuation across the entire expression range (Fig. 8 G). Instead, *C_L_* scaled linearly with *C_total_*. Consistent with their time-lapse kinetic profiles (Figs. 4 and 5), where ectopic assemblies still form despite tens-of-fold increases over wild-type levels, this linear scaling reflects inefficient thermodynamic partitioning into a distinct dense phase. Consequently, these assemblies capture only a minor fraction of the overexpressed pool, as exemplified by Sla1 reaching a nearly ideal 1:1 trajectory, leaving most of the protein in the bulk cytoplasm.

### Pan1’s buffering threshold (***C****_onset_*) is actively modulated by opposing kinase (Akl1) and phosphatase (Scd5-Glc7) activities *in vivo*

Phosphorylation of Pan1 and several key adaptor proteins by Ark1/Prk1/Akl1-family protein kinases triggers CME coat disassembly (Bourgoint et al., 2018; Huang et al., 2003; Roelants et al., 2017; Sekiya-Kawasaki et al., 2003; Toshima et al., 2005), whereas Scd5 recruits the PP1 phosphatase Glc7 to CME sites to dephosphorylate Pan1 and its binding partners (Chang et al., 2002; Chi et al., 2012; Henry et al., 2002; Zeng et al., 2007). In line with these opposing kinase-phosphatase activities, our screen revealed that Pan1, Scd5, and Akl1 exhibit concentration-buffering behavior (Fig. 8 G). To investigate whether Pan1’s *C_onset_*is governed by its phosphorylation state, we overexpressed Akl1 or Scd5 in cells expressing Pan1 at endogenous levels.

Upon Akl1 overexpression, Pan1 failed to form cortical CME foci and instead appeared diffuse throughout the cytoplasm (Fig. 9 A), indicating that Akl1 kinase activity inhibits Pan1 assembly at CME sites. Akl1 induction for 180 min resulted in a 68.7% increase in mean *C_L_* relative to the uninduced baseline (*p* = 0.0001, Mann-Whitney test). These results demonstrate that excess kinase activity prevents Pan1 buffering at wild-type concentration thresholds, effectively elevating its operational phase boundary.

**Figure 9.**
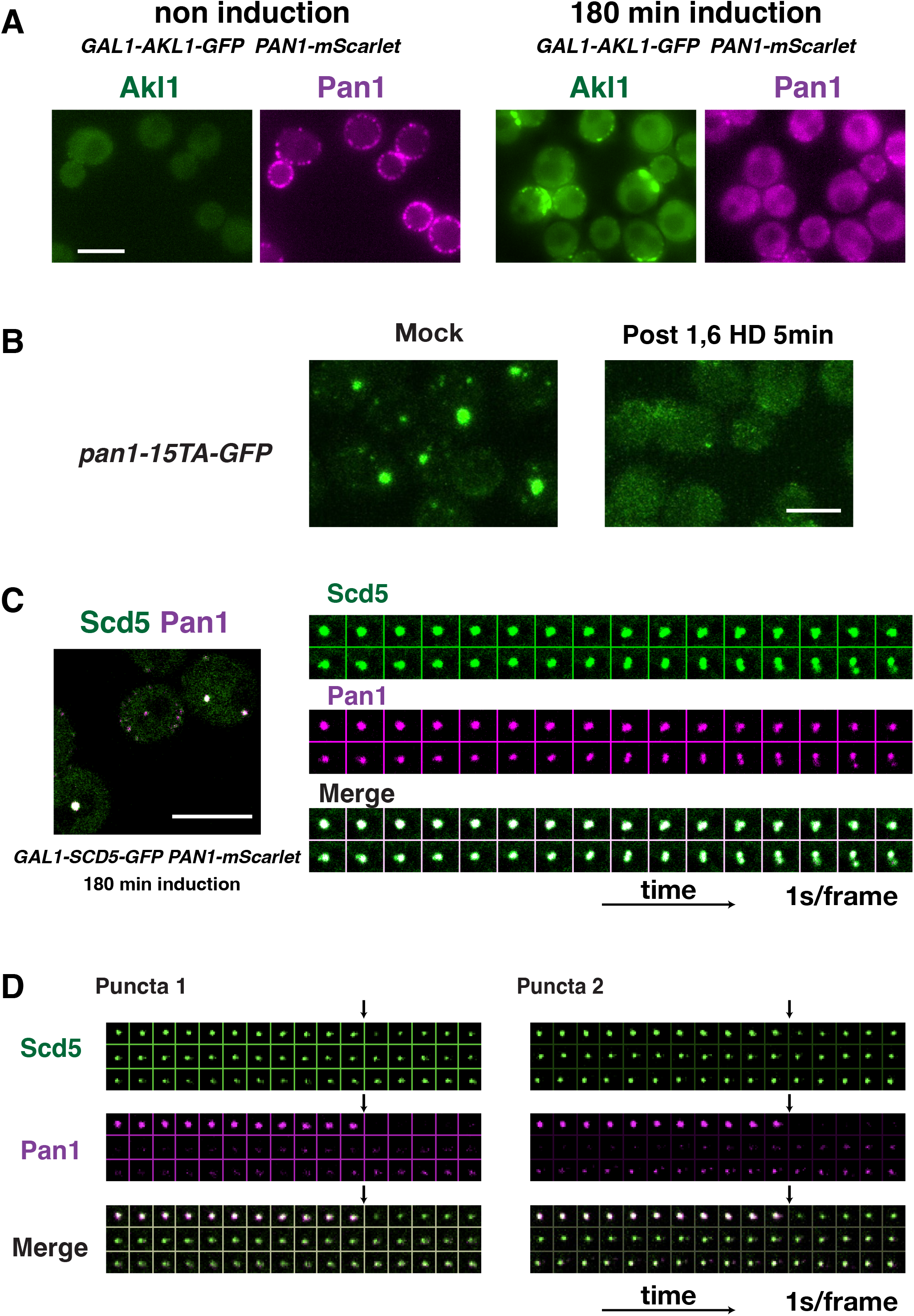
The balance between kinase (Akl1) and phosphatase (Scd5-Glc7) activities modulates *C_onset_* for endogenously expressed Pan1. **(A)** Two-color live-cell imaging of cells expressing *GAL1*-driven Akl1-GFP and endogenously expressed Pan1-mScarlet before (left) and after 180 min of induction (right). **(B)** MIPs of z-stack live-cell imaging of cells expressing *pan1-15TA-GFP* untreated (left) or treated (right) with 10% 1,6-HD for 5 min. **(C)** Live-cell imaging (left) and representative time-lapse montage (right) of cells expressing *GAL1*-driven Scd5-GFP and endogenous Pan1-mScarlet following 180 min induction. Note that Scd5 and Pan1 colocalized at cytoplasmic puncta that deform/split concurrently, presumably driven by the mechanical force generated by assembling actin (see Fig. 7 B). **(D)** Representative time-lapse FRAP montages of individual cytoplasmic puncta of cells expressing *GAL1*-driven Scd5-GFP and endogenous Pan1-mScarlet following induction. Arrows indicate the photobleaching event. Scale bars, 5 *μ*m

To further confirm that direct phosphorylation of Pan1 governs its phase behavior, we examined a phosphorylation-resistant mutant *pan1-15TA-GFP,* expressed from its endogenous locus, in which 15 of the 18 known consensus Akl1/Ark1/Prk1 kinase motifs are mutated from threonine to alanine (Toshima et al., 2005). In these mutant cells, preventing phosphorylation enhanced Pan1’s intrinsic assembly propensity, causing pan1-15TA-GFP expressed at endogenous levels to form cytoplasmic assemblies that were sensitive to 1,6-hexanediol, a hallmark of assemblies driven by weak hydrophobic interactions (Fig. 9 B).

Conversely, when the phosphatase-targeting factor Scd5 was overexpressed, two-color imaging showed that Pan1 colocalized with dynamic cytoplasmic Scd5 assemblies (Fig. 9 C). We hypothesize that formation of these Scd5-Pan1 assemblies is driven by Scd5-mediated Glc7 dephosphorylation of Pan1. To test this hypothesis, we performed dual-color FRAP with simultaneous photobleaching of both fluorophores in these assemblies. Because these structures are physically motile, we restricted our analysis to assemblies that remained within the focal plane throughout time-lapse imaging. Representative recovery traces reveal that Scd5 fluorescence recovered substantially faster than co-localized Pan1. Strikingly, despite complete photobleaching, Scd5 displayed pronounced fluorescence intensity in the first post-bleach frame due to rapid exchange during the bleaching pulse itself. This kinetic decoupling strongly supports a model in which Scd5 acts as a catalytic trigger, targeting Glc7 dephosphorylation of Pan1 and promoting subsequent Pan1 phase separation. Consistently, 180 min of Scd5 expression induction resulted in a 34.6% reduction in mean *C_L_* compared to 0 min (*p* = 0.0002, Mann-Whitney test), indicating that excess Glc7 phosphatase activity lowers the operational buffering onsets to trigger Pan1 assembly.

Together, our results demonstrate that altering the balance between kinase (Akl1) and phosphatase (Scd5-Glc7) activities dynamically modulates *C_onset_* for endogenously expressed Pan1.

## Discussion

By systematically evaluating biophysical protein-assembly properties and mapping phase behavior across the core yeast endocytic proteome *in vivo*, we established a quantitative framework to identify CME proteins that can assemble into enlarged structures with distinct biophysical properties in a crowded cellular environment. These findings demonstrate that along with Ede1, the seven newly identified candidates form an interconnected condensation network of highly conserved kinases, phosphatase recruiters, and their substrates. Reversible phosphorylation dynamically tunes phase boundaries (*C_onset_*) to choreograph CME protein assembly and turnover, a mechanism likely to represent a general principle applicable to endocytosis in diverse eukaryotes.

### An *in vivo* quantitative screen uncovers a hierarchical network of CME scaffolds and clients

By profiling 21 core CME components with our *in vivo* dose-response platform and expanding on prior Ede1 studies by explicitly quantifying its operational buffering onset (*C_onset_*), we generated a comprehensive biophysical atlas spanning the endocytic timeline. Based on these quantitative parameters (summarized in Table S1), the endocytic landscape partitions into three distinct functional classes: primary scaffolds, low-affinity homotypic assemblers, and heterotypic clients (Fig. 10 A).

**Figure 10.**
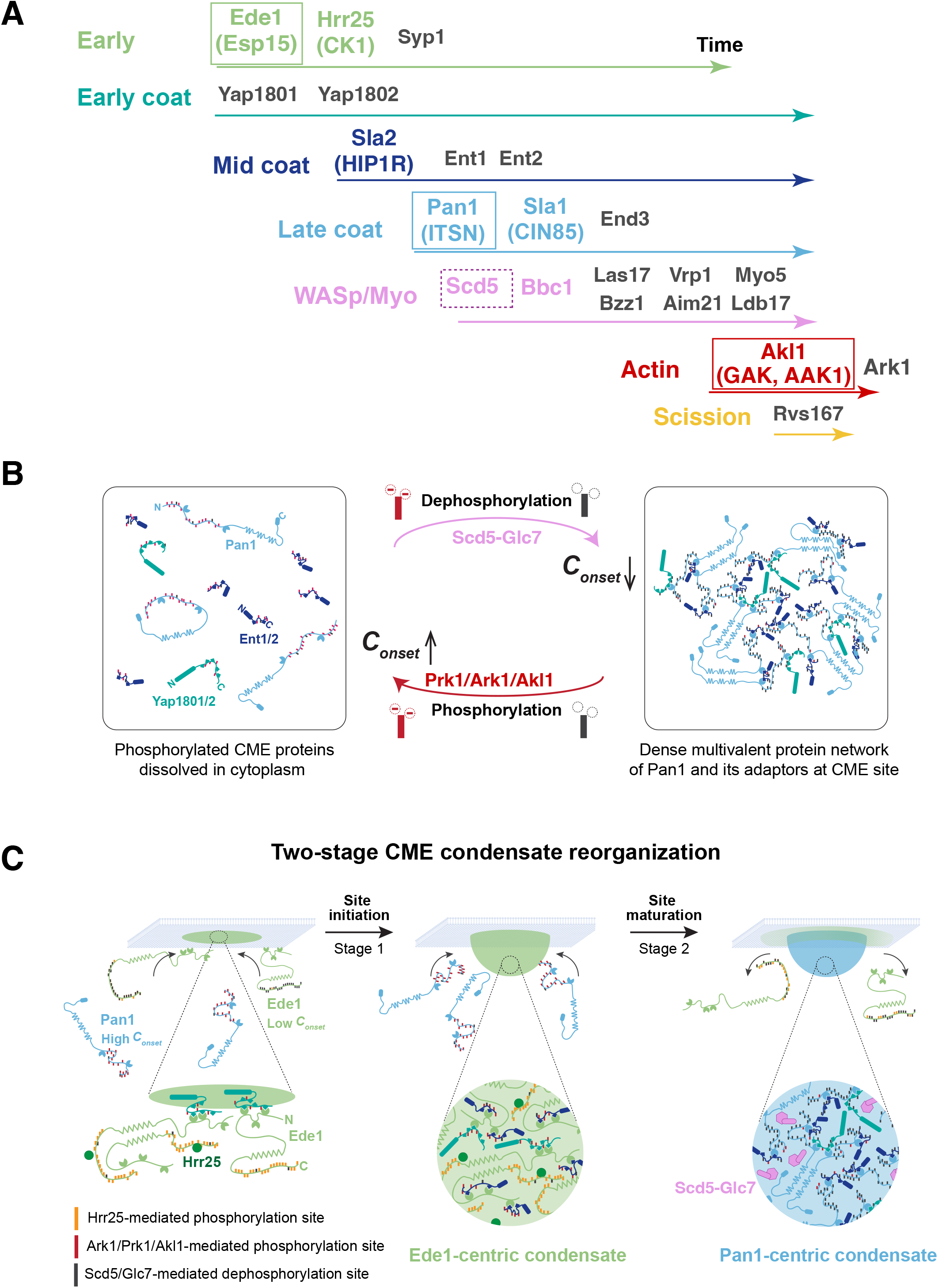
Phosphoregulation drives hierarchical assembly and two-stage reorganization of the CME condensate network. **(A)** A hierarchical condensate network of CME scaffolds and clients. A timeline illustrating the sequential recruitment of 22 core proteins to a CME site. Proteins above each arrowed line belong to the same module and share a similar lifetime, represented by line length, with start and end points marking their recruitment and disassembly. Our results suggest that CME site assembly and turnover are driven by three distinct operational protein classes: primary condensate scaffolds (boxed; Ede1, Pan1, Akl1), low-affinity homotypic condensate assemblers (colored text; Hrr25, Sla2, Sla1, Bbc1), and heterotypic clients (black text; 14 CME proteins that do not form apparent concentration-dependent assemblies). Scd5 functions as a catalytic co-scaffold (dashed box). Mammalian orthologs in parentheses. **(B)** Reversible assembly of the multivalent Pan1-adaptor network tuned by a phosphorylation dial. (Left) Ark1/Prk1/Akl1-mediated phosphorylation (red sticks) of Pan1 and adaptor IDRs introduces electrostatic repulsion that destabilizes multivalent EH-NPF networking, driving coat disassembly. This phosphorylation elevates Pan1’s *C_onset_* above cytoplasmic concentrations, releasing CME proteins into the dilute cytoplasm. Pan1 contains two N-terminal EH domains flanked by 18 phosphorylation sites within its IDR, central coiled-coil regions for oligomerization, and a C-terminal PRD. Ent1/2 and Yap1801/2 feature N-terminal ENTH/ANTH domains and C-terminal IDRs enriched in NPF motifs interspersed with 3 to 5 phosphorylation sites. (Middle) A reversible phosphorylation dial controls the transition between the solubilized cytoplasmic pool and the condensed network. Scd5-directed Glc7 dephosphorylation removes negative charges (⊖) from IDR linkers, lowering Pan1’s *C_onset_* and promoting dense network assembly, whereas Prk1/Ark1-mediated phosphorylation restores negative charges to drive disassembly. (Right) During late coat assembly, Scd5-directed Glc7 dephosphorylates multiple sites within IDRs of Pan1 and adaptors (gray sticks). Neutralizing these negative charges relieves electrostatic repulsion, thereby lowering Pan1’s *C_onset_*and enhancing multivalent EH-NPF avidity. Together with Pan1 oligomerization, this enhanced avidity triggers dense network assembly prior to actin polymerization. Other Pan1 binding partners (e.g., Sla1, Sla2, End3) are omitted for clarity. **(C)** Proposed model for two-stage condensate reorganization from initiation to maturation during CME. Stage 1 (Site Initiation): Early adaptors recruit Ede1 to the plasma membrane, where 2D spatial confinement allows Ede1 to exceed its low threshold concentration *(C_onset_)* and form an Ede1-centric condensate. Ede1 contains three N-terminal EH domains and a central coiled-coil region for oligomerization. Hrr25 phosphorylates multiple sites within Ede1’s C-terminal IDRs (orange sticks) to maintain condensate fluidity and dynamic exchange, presumably preventing gelation due to Ede1’s low *C_onset_*. Pan1’s high baseline *C_onset_* prevents its premature phase separation. Concentrated adaptors within the Ede1-centric condensate cooperatively engage Pan1 and recruit the Scd5–Glc7 phosphatase complex. Stage 2 (Coat Maturation): Scd5-Glc7 dephosphorylates Pan1 and associated adaptors (gray sticks), lowering Pan1’s *C_onset_* to trigger dense Pan1-centric condensate assembly. Enhanced multivalent binding avidity causes Pan1 to outcompete Ede1 for shared adaptors, driving Ede1 displacement prior to actin machinery assembly. Note: Additional Pan1 binding partners and downstream steps (actin-mediated invagination, scission, and site disassembly) are omitted. Domain architectures and color schemes for Pan1, Yap1801/2, and Ent1/2 are defined in **(B)**.

The primary scaffolds Ede1 (early module), Pan1 (mid/late coat), and Akl1 (actin module) act as stage-specific network nodes whose properties are well-suited to drive sequential CME stages (Fig. 10 A). Each of these three proteins displays robust concentration buffering marked by distinct transition points in their titration plots (Figs. 3 F, 4 C, 5 C, and 8 G). While yeast Pan1 oligomerizes via its central coiled-coil domains (Miliaras et al., 2004; Pierce et al., 2013), plant Pan1 (AtEH1) undergoes phase separation, a property conserved in chimeric constructs containing yeast Pan1’s disordered domain (Dragwidge et al., 2024). Akl1 is a protein scaffold that partitions into both spherical puncta and rigid cables upon overexpression (Figs. 5 C, 6 D, and 8 F). In contrast, its close homolog Ark1 failed to form such structures (Fig. S2 K), likely because Akl1 possesses a significantly larger C-terminal IDR containing a predicted prion-like domain (Fig. 1 B). Distinct from these structural scaffolds, Scd5 functions as a catalytic co-scaffold. Although Scd5 displays concentration buffering (Fig. 8 G), its ultra-fast exchange (Fig. 8 E) and kinetic decoupling from Pan1 (Fig. 9 D) suggest that Scd5 acts by recruiting Glc7 to dynamically lower Pan1’s *C_onset_*and trigger localized co-assembly.

In contrast, low-affinity homotypic assemblers Hrr25 (early module), Sla2 and Sla1 (mid/late coat), and Bbc1 (WASp/Myo module) exhibit weak intrinsic self-affinity and lack concentration buffering within physiological ranges (Fig. 8 G). However, high-level overexpression forces them to form distinct puncta (Figs. 4 and 5), revealing a latent capacity to self-assemble when protein density is artificially elevated. These findings align with prior work: Hrr25 was proposed to act as a client in cytoplasmic P-bodies (Zhang et al., 2016), while Sla2 phase separates *in vitro* (George Draper-Barr et al., 2025). Sla1 is an extreme case: despite undergoing thermoresponsive phase transitions under specific *in vitro* perturbations (Bergeron-Sandoval et al., 2021), its linear, unbuffered titration indicates that Sla1 lacks partitioning homeostasis and deviates from canonical LLPS *in vivo* (Fig. 8 G). Finally, while Bbc1 was not previously linked to condensates, its SH3 domain, PRDs, and large IDR (Fig. 1 B) provide the structural basis for this self-assembly.

Finally, the remaining 14 CME components that did not form enlarged assemblies in our screen (Fig. S2 and Table S1) constitute the third class: heterotypic clients that do not undergo autonomous self-association (Fig. 10 A). These heterotypic clients include early coat proteins (Yap1801/2 and Syp1(yeast FCho)), mid-and late-coat proteins (Ent1/2 and End3), and actin assembly regulators (Las17 and Myo5). Rather than driving assembly independently, these well-conserved core CME proteins rely on multivalent heterotypic interactions (such as EH-NPF, coiled-coil, and SH3-PRD pairings) to interact with the primary scaffolds, Ede1 and/or Pan1 (Owen et al., 2025). Suppressing autonomous self-affinity in these heterotypic clients, such as PLD-containing adaptors (Yap1801/2, Ent1/2), serves as a crucial evolutionary safeguard against ectopic nucleation, consistent with recent work on mammalian Epsin1 (Sarkar et al., 2025).

### Multisite phosphorylation in IDRs dynamically tunes CME scaffold material states and phase behavior

Here, we demonstrate that Pan1’s *C_onset_* is dynamically governed by balanced kinase and phosphatase activities (Fig. 9). Ark1/Prk1/Akl1 kinases target the [*L*/*I*]*xxQxTG* consensus motif present across multivalent IDRs of the CME machinery, clustering most densely on scaffold protein Pan1 (18 motifs flanking its EH domains), alongside clients including early adaptors Yap1801/2 (3-4 motifs interspaced with 5 NPF motifs) and mid-adaptors Ent1/2 (5 motifs separating 2 NPF motifs) (Huang et al., 2003; Miao et al., 2018). Given that Pan1’s EH domains canonically bind NPF motifs of Yap1801/2 and Ent1/2 (Maldonado-Baez et al., 2008; Wendland and Emr, 1998), Ark1/Prk1/Akl1-mediated multisite phosphorylation likely introduces steric and electrostatic repulsions at adjacent IDR motifs to disrupt multivalent EH–NPF avidity (Bah and Forman-Kay, 2016), elevating Pan1’s *C_onset_* to trigger coat disassembly at the final stage of CME. We propose that following coat disassembly, phosphorylated Pan1 and these adaptors are recruited to new CME sites established by Ede1, where Scd5-directed Glc7 locally dephosphorylates multiple sites within IDRs (Chi et al., 2012; Zeng et al., 2007) to reduce electrostatic repulsions, promote multivalent engagement, and lower Pan1’s *C_onset_* to prime coat condensation (Fig. 10 B).

The *pan1-15TA* mutant indicates that dephosphorylating Pan1 alone is sufficient to trigger condensation (Fig. 9 B). However, while dephosphorylating Pan1 alone leaves assemblies largely 1,6-HD sensitive (Fig. 9 B), network-wide CME dephosphorylation via Scd5 overexpression lowers Pan1’s *C_onset_* and strengthens scaffold integrity, conferring ∼45% 1,6-HD resistance (Fig. 5 A). Furthermore, although point mutations in NPF-binding pockets of all Pan1 EH domains yield only mild phenotypes (Suzuki et al., 2012), deleting EH domains with their flanking IDRs causes lethality (Bradford et al., 2015), suggesting that multivalent assembly platforms, rather than isolated EH-NPF interactions, are essential for Pan1 function. Thus, while Pan1 dephosphorylation alone is sufficient to initiate coat condensation, multisite dephosphorylation across IDRs of both scaffold (Pan1) and client adaptors (Yap1801/2, Ent1/2) appears necessary to establish a dense multivalent network to stabilize the CME coat prior to actin assembly.

Like Pan1, Ede1 harbors N-terminal EH domains that interact with NPF motifs in Yap1801/2 and Ent1/2 (Aguilar et al., 2003; Howard et al., 2002). However, Ede1 possesses a notably low *C_onset_* and carries only one [*L*/*I*]*xxQxTG* consensus motif (Huang et al., 2003). Instead, more than 20 Ser/Thr sites clustered within its C-terminal IDR are phosphorylated by Hrr25 (Fig. 10 A). Hrr25-mediated Ede1 phosphorylation promotes efficient CME initiation (Peng et al., 2015). Our FRAP analysis reveals moderate exchange kinetics for Hrr25 within assemblies upon overexpression (t_1/2_ = 23.4 s) (Fig. 8 A), suggesting a sustained presence to support continuous Ede1 phosphorylation. By preserving Ede1 condensate fluidity and preventing gelation due to its low *C_onset_*, Hrr25 phosphorylation appears to fine-tune scaffold interactions to ensure functional CME site initiation.

We propose that because high-affinity “lock-and-key” interactions risk kinetic traps that impede disassembly, multivalent, low-affinity IDR interactions regulated by reversible phosphorylation are utilized to allow the CME machinery to achieve rapid assembly and swift turnover. Governed by stage-specific protein kinases and phosphatases (Hrr25, Scd5-Glc7, and Ark1/Prk1/Akl1), multisite IDR phosphorylation provides a versatile biophysical dial to modulate *C_onset_* , scaffold material properties, and condensate dynamics across the CME pathway, illustrating a broader regulatory paradigm in active phase separation (Soding et al., 2020).

### Differential condensation thresholds for Ede1 and Pan1 underlie a two-stage CME network reorganization

While Ede1 and Pan1 share similar domain architectures, comparable wild-type cytoplasmic concentrations (∼140 nM) (Boeke et al., 2014; Gagny et al., 2000), and shared interaction partners (Aguilar et al., 2003; Boeke et al., 2014; Howard et al., 2002; Toshima et al., 2007; Wendland and Emr, 1998), Ede1 arrives at CME sites tens of seconds before Pan1 and disassembles right before Pan1 recruits the actin assembly machinery to drive invagination (Carroll et al., 2012; Sun et al., 2024b). Our quantitative *in vivo* measurements provide a biophysical rationale for why variable Ede1 initiation precedes the strictly timed recruitment of Pan1 (Fig. 8 G). Ede1’s *C_onset_* (∼ 284 nM) is within two-fold of its native cytoplasmic concentration, enabling local concentration on the plasma membrane to readily push it across the phase boundary. Conversely, Pan1 exhibits a high *C_onset_* (∼ 1357 nM), nearly ten-fold above its native cytoplasmic concentration, reflecting a steep, kinase-regulated repressive barrier that is abolished in the phosphorylation-deficient *pan1-15TA* mutant (Fig. 9 B).

Building on key discoveries from our analysis of Ede1 and seven new candidates, we propose a unified, two-stage condensation model that can explain the physical principles orchestrating the CME pathway progression (Fig. 10 C):

In Stage 1 (Site Initiation), early lipid-binding adaptors (such as Yap1801/2 and Syp1) recruit Ede1 to the plasma membrane (Sun et al., 2024b). 2D spatial confinement allows Ede1 to exceed its low *C_onset_*and drive Ede1-centric condensate formation. Although Pan1 can also interact with early adaptors, its baseline phosphorylation maintains a high *C_onset_*that prevents premature phase separation, ensuring that Ede1 initiates condensation exclusively. The protein kinase Hrr25 associates with Ede1, maintaining its proper dynamic state via sustained phosphorylation (Peng et al., 2015). Ede1-centric condensates act as functional reaction hubs that accelerate recruitment kinetics of mid-arriving coat proteins (such as Ent1/2 and Sla2), which also carry lipid-binding domains (Aguilar et al., 2003; Boeke et al., 2014; George Draper-Barr, 2025; Howard et al., 2002). Then, by vastly increasing local binding site density, the concentrated adaptors cooperatively engage Pan1 (Aguilar et al., 2003; Boeke et al., 2014; Howard et al., 2002; Maldonado-Baez et al., 2008; Toshima et al., 2007; Wendland and Emr, 1998) and recruit the Scd5-Glc7 phosphatase complex (Zeng et al., 2007).

In Stage 2 (Coat Maturation & Force Generation), Scd5 targets Glc7 to the CME site, where it dephosphorylates Pan1 and its binding partners, such as Yap1801/2, Ent1/2, and Sla1, across a few dozen [*L*/*I*]*xxQxTG* consensus motifs within their IDRs (Chi et al., 2012; Zeng et al., 2007). Dephosphorylation of Pan1 and its partners lowers Pan1’s *C_onset_* to trigger phase separation and increase multivalent interaction density, consolidating partners into a Pan1-centric condensate. Meanwhile, sustained Hrr25 activity maintains Ede1 phosphorylation and condensate fluidity during network expansion. By selectively enhancing Pan1’s binding avidity toward itself and its binding partners, Scd5-Glc7-mediated dephosphorylation shifts the binding equilibrium so Pan1 outcompetes Ede1 for shared adaptors, driving Ede1’s spatial displacement (Mund et al., 2018) and subsequent disassembly alongside Hrr25 prior to actin assembly initiation (Peng et al., 2015; Stimpson et al., 2009). Assisted by Sla1 and Bbc1, the mature Pan1-centric condensate recruits Las17 (yeast WASp) and Myo5 through an interconnected SH3-PRD multivalent network. Echoing how vertebrate WASp condensation drives actin nucleation (Case et al., 2019), this condensed multivalent network triggers an explosive, synchronized burst of Arp2/3-dependent actin assembly (Sun et al., 2017). This Pan1-driven scaffold enables Sla2 and Ent1/2 to anchor actin filaments to the membrane, transmitting mechanical forces against high turgor pressure to drive membrane invagination (Skruzny et al., 2012). Finally, Ark1/Prk1 family kinases are recruited to the CME site through indirect interactions with actin filaments (Cope et al., 1999; Fazi et al., 2002), phosphorylating components within the Pan1-centric condensate to raise Pan1’s *C_onset_*above cytosolic levels and drive condensate dissolution.

This two-stage condensation model reveals an elegant evolutionary adaptation. Distinct phosphorylation circuits targeting divergent IDRs in Ede1 and Pan1 as well as their shared adaptors precisely modulate phase boundaries and recruitment kinetics. This Ede1-to-Pan1 transition ensures that robust site initiation is followed by the formation of a dense Pan1-centric network that triggers explosive actin assembly and rapid subsequent disassembly. Such spatiotemporally regulated condensate assembly can overcome biophysical constraints, such as cytosolic noise, 3D diffusion limits, and mechanical slippage, and functionally decouple site initiation from force generation. Consistently, acute Pan1 depletion leaves Ede1 initiation intact (Bradford et al., 2015; Sun et al., 2015), whereas peroxisome-targeted Pan1 bypasses Ede1 entirely to drive actin assembly (Enshoji et al., 2022). Strikingly, cells lacking most early factors (including Ede1) remain viable and undergo CME, albeit less efficiently (Brach et al., 2014). In contrast, Pan1 and Scd5 are indispensable for viability, underscoring their roles in forming essential commitment nodes that license coat maturation.

### Limitations, capabilities, and broader utility of the *in vivo* dose-response strategy

Our high-throughput *in vivo* dose-response assay involves technical trade-offs. Because diffraction-limited imaging cannot resolve sub-microscopic oligomers, their early accumulation manifests as a minor increase in *C_L_*before they grow into detectable assemblies. Terminal fluorescent protein tag placement can also alter protein conformation or obscure interaction surfaces (Snapp, 2005). Indeed, while both engineered Sla1-fluorescent protein fusions tested here are endogenously functional, only N-terminally tagged Sla1 formed visible puncta upon overexpression (Fig. 4 D and Fig. S2 Q), likely because C-terminal tagging alters multivalent interaction kinetics. These nuances highlight how false negatives and positives may have arisen in our screen. Nevertheless, our quantitative analysis of Ede1 and seven newly identified candidates takes a crucial step toward revealing key principles governing sequential CME protein recruitment. Future platform iterations could implement multi-locus induction to co-express multi-subunit complexes at natural stoichiometric ratios, while pairing minimal tags (such as SNAP/CLIP or split-GFP) with FLIM to enable real-time micro-viscosity measurements inside condensates.

Our pipeline leverages *in vivo* single-cell titrations to extract threshold concentrations (*C_onset_*), providing a quantitative framework for *in vitro* reconstitution to test our two-stage condensation model. Additionally, this dataset provides a benchmark for training predictive algorithms to classify scaffolds versus clients. Beyond endocytosis, this dose-response framework offers a general strategy to map phase boundaries across diverse multivalent assemblies *in vivo*.

## Methods and Materials

### Strains and cell culture conditions

Yeast strains were grown in standard rich media (YPD) or synthetic media (SD) supplemented with the appropriate amino acids. The yeast strains used in this study are listed in Table S2. The Gal1 promoter was endogenously integrated at the 5’ end of each open reading frame (Longtine et al., 1998). For induction experiments, yeast cells were grown to mid-log phase at 25°C and Gal1 promoter activation was induced by addition of 50 nM β-estradiol (Sigma-Aldrich, E2758). Chemical treatments were for 5 min for 10% 1,6-hexanediol (Sigma-Aldrich, 240117) or for 5-10 min for 250 µM CK-666 (Sigma-Aldrich, SML0006).

### Western blotting

5 *OD*_600_ of yeast were harvested at various time points after induction. Cell pellets were resuspended in 20% (w/v) trichloroacetic acid (TCA) and snap-frozen in liquid nitrogen. After thawing, precipitated proteins were pelleted by centrifugation, neutralized, and solubilized in SDS–PAGE sample buffer. Proteins were resolved on 4-20% Tris-glycine SDS–polyacrylamide gradient gels (37.5:1 acrylamide:bis) and transferred to nitrocellulose membranes by wet transfer using a Mini Trans-Blot cell (Bio-Rad) in Tris-glycine transfer buffer containing 20% (v/v) methanol, overnight at 30 V at 4 °C.

Nitrocellulose membranes were first blocked for 1 hour at room temperature in Tris-buffered saline (TBS; 20 mM Tris-HCl pH 7.5, 150 mM NaCl) containing 5% (w/v) nonfat dry milk and 0.2% (v/v) Tween 20. Membranes were then incubated for 1 hour at room temperature with primary antibodies, mouse anti-GFP (1:1,000) from Roche and mouse anti-Pgk1 (1:40,000; loading control) from Invitrogen, diluted in TBS with 0.2% Tween 20 (TBST) plus 5% milk (TBS-T). Membranes were washed three times for 5 min in TBS-T, incubated for 1 hour with a donkey anti-mouse fluorescent secondary antibody (1:20,000) from LI-COR Biosciences, washed three more times for 5 min in TBST, and rinsed once in TBS. Blots were imaged on an Odyssey CLx scanner (LI-COR Biosciences), and single band intensities were quantified in Image Studio (LI-COR). The signal for the GFP-tagged protein of interest was normalized to Pgk1 in the same lane. Final band intensities for proteins were calculated as fold changes relative to wild type.

### Live-cell imaging

#### Confocal imaging

Cells were immobilized in concanavalin A-coated 8-well imaging chambers prior to imaging. Confocal images were acquired using a Nikon Eclipse Ti2 AX R confocal microscope controlled by NIS-Elements software and equipped with a Plan Apo λD 100× oil-immersion objective, NA 1.45, using Type F immersion oil. GFP-tagged proteins were excited using the 488 nm laser. Laser power and detector gain were adjusted within the linear range for each yeast strain and held constant across analyses. Across experiments, 488 nm laser power ranged from 1.5% to 15%, with detector gain adjusted as needed to avoid saturation. Gain ranged from 1.5 to 10.

Images were acquired using resonant bidirectional resonant scanning with 2x line averaging, a 1024 × 1024-pixel frame size, and a pinhole size of approximately 120.9 µm, corresponding to approximately 2.5 Airy units. Z-stacks were collected using the Nikon AX Piezo Z drive and contained 23 optical sections with a step size of approximately 0.30 µm and 165 ms exposure per section.

#### Confocal imaging with NSPARC (Nikon Spatial Array Confocal) detector

Cells were immobilized in concanavalin A-coated 8-well imaging chambers prior to imaging. NSPARC (Nikon Spatial Array Confocal) confocal imaging was performed using a Nikon Eclipse Ti2 microscope equipped with a 100×1.4 NA Plan Apo D oil objective. To investigate the protein recruitment order for formation of enlarged concentration-dependent assemblies relative to the endocytic actin marker, yeast strains co-expressing GFP-tagged assembly-forming proteins and 2x mScarlet-tagged Sac6 were tracked via high-speed two-color movie acquisition (0.5sec/frame) after 180 min induction. Dual-channel image collection was mediated by a high-speed Resonant scanner operating in bidirectional mode. Channels were acquired sequentially via line-by-line switching, alternating the 488-nm and 561-nm excitation lines at each scan line. Spatial crosstalk and channel bleed-through were empirically evaluated using single-labeled control strains and determined to be negligible.

#### Fluorescence recovery after photobleaching

Fluorescence recovery after photobleaching (FRAP) was performed on enlarged concentration-dependent assemblies to evaluate their dynamic material properties *in vivo*. NSPARC confocal imaging was performed using a Nikon Eclipse Ti2 microscope equipped with a 100x 1.4 NA Plan Apo D oil objective. Individual puncta were targeted as regions of interest (ROIs) alongside separate unbleached control puncta and extracellular background references. Time-lapse sequences were initiated by capturing 10 pre-bleach baseline frames (1sec/frame). Active photobleaching was subsequently performed using a focused 488-nm laser at 90-100% power for 2 s. The laser raster pattern was controlled via a high-precision Galvano scanner configured in bidirectional acquisition mode with a precise pixel dwell time of 2 µs per spot. This setup ensured a highly efficient, continuous, and deeply reduced fluorescence intensity within the targeted macromolecular structures. Following the photobleaching loop, post-bleach fluorescence recovery was recorded by acquiring images at regular 1s intervals for a total monitoring window of 1-2min using highly attenuated laser power to prevent unintended imaging-induced photobleaching. Raw NSPARC datasets were computationally reconstructed and processed utilizing Nikon NIS-Elements software.

### Image Analysis

#### Image segmentation and quantification

Image analysis was automated using a custom pipeline implemented in Python, which builds upon our previously described automated analysis workflow (Sun et al., 2024b). ImageJ (imagej.nih.gov/ij/) functions were used and ported to Python with PyImageJ (Rueden et al., 2022). The analysis code is available at github.com/geanhu/cme-analysis-2025.

The differential inference contrast (DIC) channel of acquired z-stacks was used as the input for cell segmentation. Background subtraction was performed using the rolling ball method with a radius of 70 pixels approximated with a paraboloid function, as implemented in ImageJ. The maximum intensity projection of the stack was then calculated, excluding the first 3 and last 3 sections (of 23 sections total), as these sections are often out of focus and extend beyond the depth of the imaged cells. A median filter with a radius of 5 pixels was then applied to reduce noise. This filtered projection was then provided as input to Cellpose (Stringer et al., 2021), a deep-learning-based segmentation model, and cells were segmented using Cellpose’s cyto model (trained only on Cellpose training data) and a cell diameter estimate of 50 pixels. Any holes internal to a single label area were filled using functions from the MorphoLibJ plugin library (Legland et al., 2016).

The GFP channel of acquired z-stacks was used as input for cytoplasmic puncta segmentation. We generated a maximum intensity projection of the stack as described above, then cropped it by 50 pixels from each edge to avoid uneven illumination at the image borders (resulting in 924 x 924 pixel images). The median intensity of all pixels that were not segmented into a cell was taken as the background value and subtracted from the image. Next, we adapted the PunctaTools method for puncta segmentation (Baggett et al., 2022). Because we observed that the fluorescence intensity of cells within a field is highly variable, puncta segmentation was performed for each segmented cell area independently. For each cell area, Laplacian of Gaussian (LoG) filters were applied using *σ* ranging from 2 pixels to 5 pixels. Then, the maximum resulting value at each pixel, normalized by *σ*^2^, was determined across the range of filter sizes. Pixels that have a value greater than X normally scaled median absolute deviations above the median were selected as potential cytoplasmic puncta centers, with the constraint that puncta centers must be at least 5 pixels apart. Puncta centers were used as seeds for watershed segmentation, where pixels with normally scaled median absolute deviations greater than Y were segmented into a puncta label. X and Y are user-defined hyperparameters that we determined for each sample by manually inspecting puncta segmentation accuracy. We discarded any resulting puncta smaller than 5 pixels as random noise mistakenly segmented.

While we used the maximum intensity projection of the GFP channel for puncta segmentation (because maximum projections tended to have higher contrast between puncta and background), the sum intensity projection of the GFP channel was used to measure intensity values (because we reasoned that sum intensity projections would correctly increase the intensity of pixels where puncta exist atop one another along the z-axis). First, we subtracted the median intensity of all pixels not segmented into cells (i.e., the background) from the entire image. To correct for cell autofluorescence, we also subtracted the average cell intensity of our wild-type strain with no fluorophore. For each cell, we determined the mean intensity of the entire cell area, the median intensity of the pixels segmented as puncta, the median intensity of the pixels not segmented as puncta (thus segmented into the cytoplasm), and the number of puncta within the cell. We reasoned that the mean is an appropriate measure for the intensity of the entire cell area since this metric should summarize the bimodal distribution of both the higher intensity puncta pixels and the lower intensity cytoplasmic pixels; in contrast, we reasoned that the median is an appropriate measure for the intensity of the puncta or cytoplasmic areas only, since these distributions would be expected to be unimodal. For each punctum, we determined the size (number of pixels segmented as puncta, converted to *m*^2^ using the pixel size set during image acquisition) and mean intensity.

For graphs of “normalized average cell intensity”, mean cell intensities at each induction time point were normalized by the mean of all the mean cell intensities measured from the POI-GFP image, thus determining the fold change over wild-type expression levels. For graphs comparing puncta sizes before and after 1,6-hexanediol treatment, we first randomly sampled an equal number of cells from the pre-treatment and post-treatment images, then compared puncta sizes within those sampled cells.

For the quantification shown in Fig. 8 G, we converted measured median intensities of the puncta (*I_D_*) to dense-phase concentration (*C_D_*), and measured median intensities of the cytoplasm (*I_L_*) to determine the dilute cytoplasm concentration (*C_L_*) , using cytosolic protein concentrations previously determined by fluorescence-correlation spectroscopy (FCS) (Boeke et al., 2014). We first determined the ratio of the average particle concentration measured by FCS to the mean wild-type cytoplasmic intensity, then multiplied the indicated intensities at 180 min post-estradiol induction by this ratio. Because Hrr25 and Akl1 were not measured in the aforementioned FCS experiments, ratios for these proteins were imputed using the ratio of Ede1 FCS cytosolic concentration to the mean of wild-type median cytoplasmic intensities of Ede1 samples imaged using the same imaging conditions and settings as for the Hrr25 and Akl1 images. For each protein, we then fit the *C_total_* versus *C_L_* data with a non-parametric LOWESS fit, using a 2/3 fraction of the data to estimate each value. The elbow of the LOWESS fit was then defined as the fit value with the largest distance normal to the line connecting the start and end of the fit.

#### Fluorescence recovery after photobleaching (FRAP) quantification

Puncta intensities, determined as described above, were normalized using the double normalization scheme defined by (Phair et al., 2004). First, at each time point, the background intensity was subtracted from both the bleached sample and the unbleached reference. Using these background-subtracted values, we calculated the amount of fluorescence loss due to photobleaching by dividing the reference pre-bleach average intensity by the reference intensity at each time point. At each time point, this value was multiplied by the bleached-sample intensity to correct for photobleaching-induced signal loss. The final relative fluorescence signal was then determined by dividing the normalized bleached-sample data by the sample’s pre-bleach average. We then fit the double-normalized FRAP data to a single-exponential model using non-linear least squares, defined as *I*(*t*) = *A*(1 − *e*^−*kt*^) + *C* where the mobile fraction is A and the recovery half-time *t*_1/2_ is determined *t*_1/2_ = *ln*(2) / *k* . Separate models were fitted for each individually bleached punctum, fits resulting in non-physical mobile fractions (*A* > 1) were excluded from analysis.

### Computational analysis of sequence features and LLPS propensity

First, through a literature search, we compiled a list of proteins that contribute key functions to CME (see Table 1, Table S1) and fetched the proteins’ amino acid sequences from the UniprotKB database.

To calculate the predicted IDR proportion for each protein, we used the Python implementation of Metapredict V3 with default parameters (Emenecker et al., 2021). We reasoned that Metapredict, a consensus predictor trained on the outputs of eight disorder predictors, would provide an approximate average snapshot of what disorder predictors in the field might produce. We used a Metapredict score threshold of 0.5 to binarize whether a single residue in a sequence is disordered (>0.5) or ordered (<0.5), then calculated the proportion of disordered residues in each protein sequence.

To calculate predicted prion-like domain proportion, we input each amino acid sequence into the command-line implementation of PLAAC (Lancaster et al., 2014), using the default *S. cerevisiae* background amino acid frequencies. Each residue was directly classified as within or not within a prion-like domain using the most likely state determined by the Viterbi algorithm, and the overall proportion of residues within a prion-like domain was then calculated for each protein.

Next, we used machine learning models to directly predict each protein’s LLPS propensity. We first used PSPHunter (Sun et al., 2024a), a recently developed model that provides per-residue LLPS predictions, and ran the command-line implementation with default parameters to determine the average predicted LLPS propensity per protein. We also used FuzDrop, a model with a comparatively simpler architecture, and directly fetched LLPS propensity values for our proteins of interest from values precomputed for the entire yeast proteome (Hardenberg et al., 2020).

Unexpectedly, a large majority of proteins we tested yielded an LLPS propensity score that approached or exceeded the binary thresholds recommended by the developers of PSPHunter and FuzDrop (0.82 and 0.61, respectively). We reasoned that PSPHunter and FuzDrop may be biased toward positive predictions for our dataset because part of their positive training sets is sourced from PhaSepDB (You et al., 2020), which includes entries of CME-related proteins previously characterized as relating to biomolecular condensate formation, such as Ede1. To circumvent this potential issue, we used an LLPS predictor that did not directly draw from PhaSepDB or other LLPS databases. To do so, we adapted FINCHES’ phase-diagram prediction function for LLPS prediction (Ginell et al., 2025; Qian et al., 2022). As FINCHES can determine relative phase diagrams comparing multiple sequences, we used the low-complexity region of the well-characterized LLPS-driving protein FUS as a positive control. To match the length of this control sequence and satisfy FINCHES’s assumption that input sequences are intrinsically disordered, for each input protein we first identified a 163-residue window that maximized the number of disordered residues. Then, phase diagrams were predicted for this input protein subsequence as compared to the FUS protein sequence, and the area under the curve (AUC) of the phase diagram for each input protein, normalized by the AUC of the phase diagram of FUS, was calculated as the final score estimating the predicted LLPS propensity for each input protein. Predictions were generated using the Mpipi-GG force field (Joseph et al., 2021) and the CALVADOS2 force field (Tesei et al., 2021), as displayed in Fig. 1 B. However, FINCHES was not originally designed for quantitative LLPS propensity prediction (Ginell et al., 2025). Additionally, FINCHES is designed to predict purely homotypic phase diagrams. In contrast, the LLPS databases used to train PSPHunter and FuzDrop contain a mix of homotypic, heterotypic, *in vivo*, and *in vitro* data, which may explain the poor agreement with FINCHES predictions.

## Declaration of Generative AI in Scientific Writing

During the preparation of this manuscript, the authors initially used Claude (Opus 4.8) and Gemini (3.5 Thinking) to help format the text structure, optimize paragraph transitions, and refine grammatical clarity. Following these automated editing suggestions, the authors rigorously reviewed, verified, and edited the output to correct grammatical errors, restore the intended meaning, and ensure scientific accuracy. We take full responsibility for the final contents of the article.

## Acknowledgements

We thank members of the Drubin/Barnes laboratory for helpful discussions. We thank Dr. Jonathan Wong, Dr. Leanna Owen, and Grace Donnelly for critically reading our manuscript and offering constructive suggestions. This work was supported by National Institutes of Health grant R35GM118149 to DGD.

