## Supplemental_figures_legends for "Phosphoregulation of hierarchical protein condensate formation during clathrin-mediated endocytosis"

### Supplemental Figure Legends

#### Supplemental Figure 1

**(A)** β-estradiol treatment does not perturb endocytic patch dynamics. Representative two-color circumferential kymographs of mother (bottom) and daughter (top) cells endogenously expressing Sla1-GFP and Sac6-2xmScarlet in the absence and presence of β-estradiol. **(B)** Maximum intensity projections (MIPs) of z-stack live-cell imaging of the indicated proteins following 180 min β-estradiol induction. Identical imaging settings and processing were used for direct comparison.

#### Supplemental Figure 2

Fourteen CME proteins failed to form enlarged puncta upon overexpression **(A-Q)**. For each protein**,** panels from left to right show: MIPs of z-stack live-cell imaging from cells endogenously expressing the indicated GFP-tagged protein, and cells expressing the *GAL1*-driven GFP-tagged protein following 180 min $\beta$-estradiol induction; Western blots as well as Pgk1-normalized quantification of protein abundance and total cellular fluorescence intensity relative to wild-type at 0 and 180 min post-induction.

### Supplemental Table Legends

#### Supplementary Table 1

Data summary of GFP- tagged Ede1 and 21 core CME proteins upon overexpression

#### Supplementary Table 2

Yeast strains used in this study.

### Supplemental Figure 1

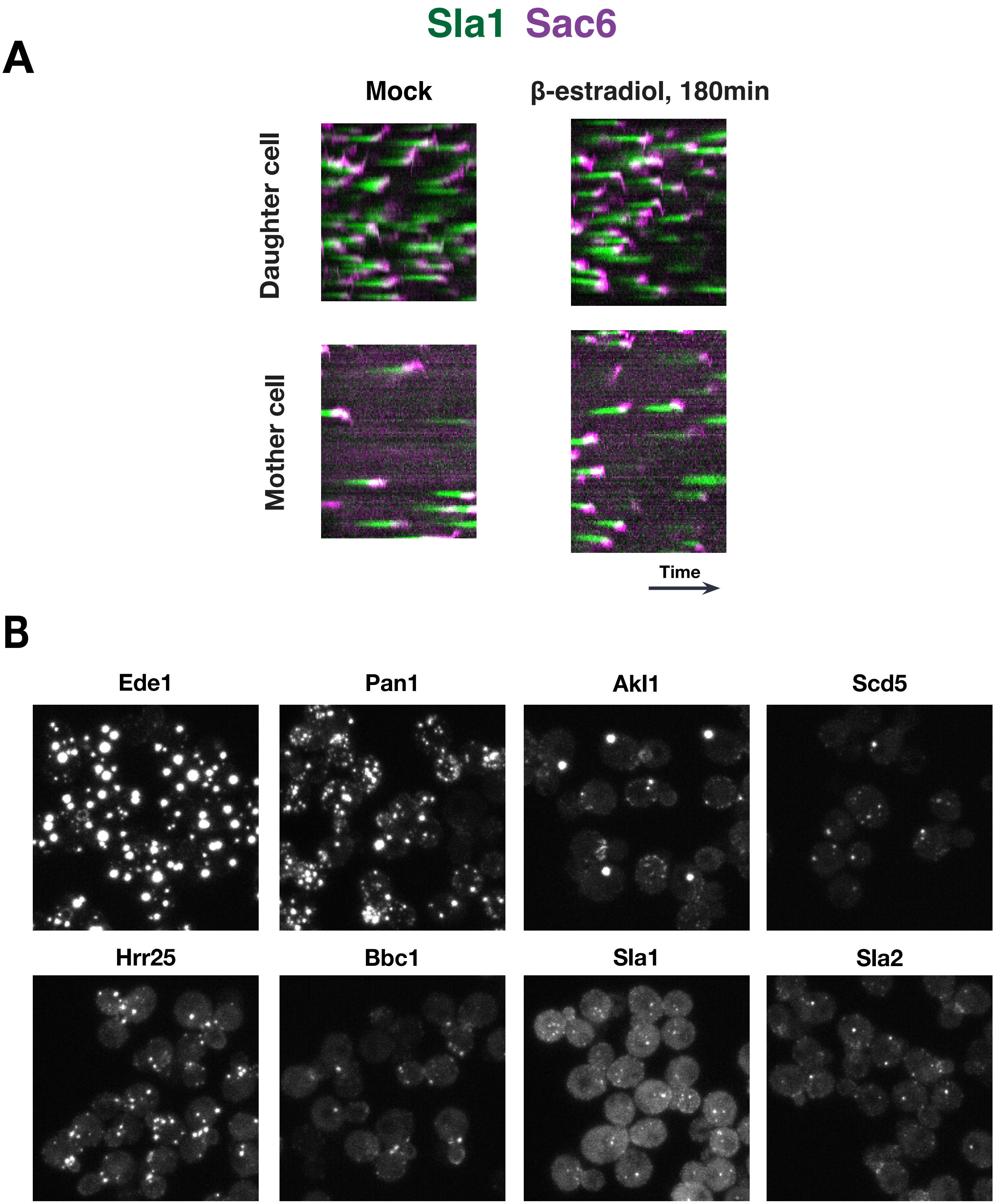

### Supplemental Figure 2_1

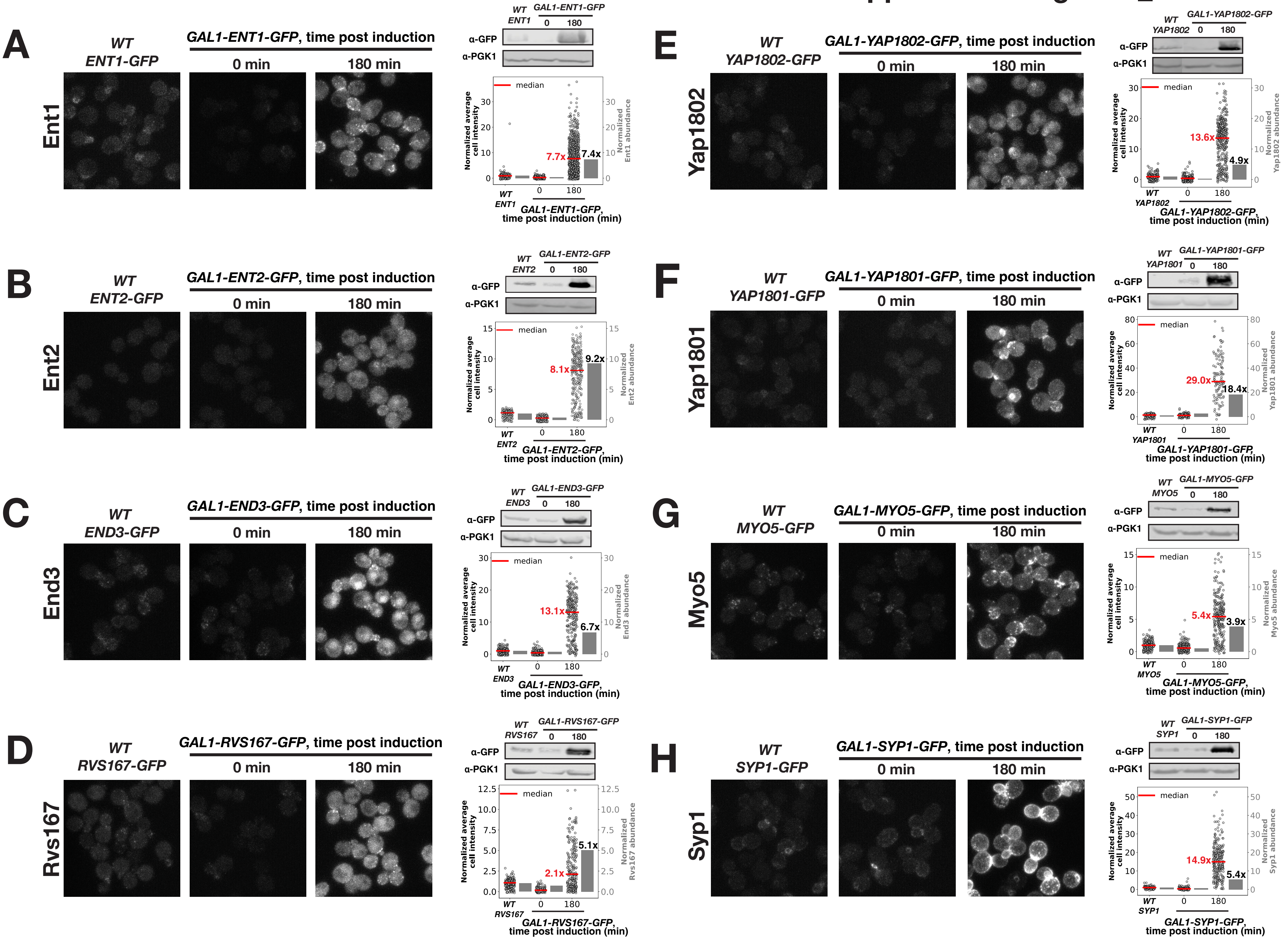

### Supplemental Figure 2_2

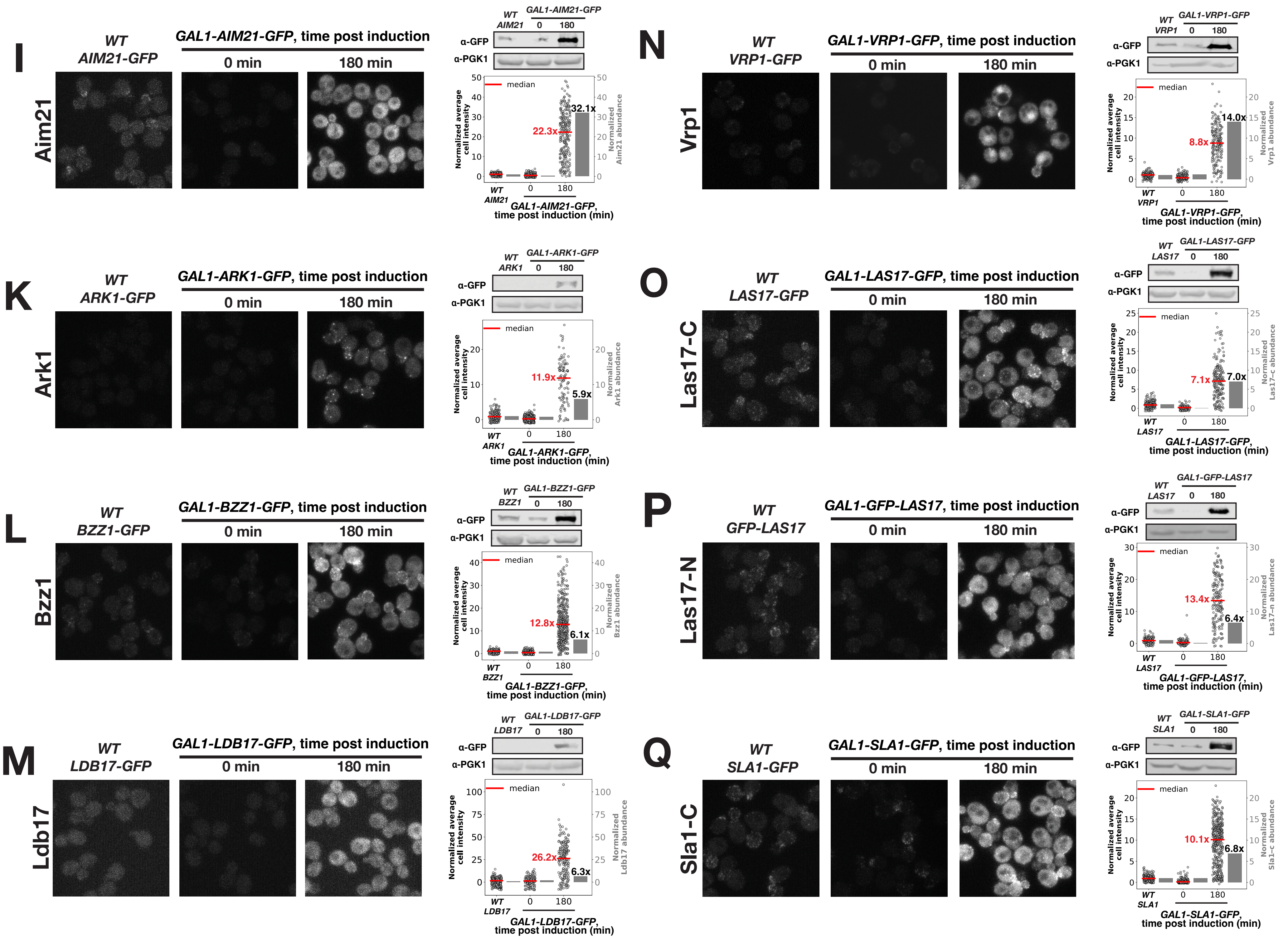
